# HYPK scaffolds the Nedd8 and LC3 proteins to initiate formation of autophagosome around polyneddylated huntingtin exon1 aggregates

**DOI:** 10.1101/780379

**Authors:** Debasish Kumar Ghosh, Akash Ranjan

## Abstract

Selective degradation of protein aggregates by autophagy is an essential homeostatic process of safeguarding cells from the effects of proteotoxicity. Among the ubiquitin-like modifier proteins, Nedd8 conjugation to misfolded proteins is prominent in stress-induced protein aggregates, albeit the function of neddylation in autophagy is unclear. Here, we report that polyneddylation functions as a post-translational modification for autophagic degradation of proteotoxic-stress induced protein aggregates. We also show that HYPK functions as an autophagy receptor in the polyneddylation-dependent aggrephagy. The scaffolding function of HYPK is facilitated by its C-terminal ubiquitin-associated domain and N-terminal tyrosine-type LC3 interacting region which bind to Nedd8 and LC3 respectively. Both Nedd8 and HYPK are positive modulators of basal and induced-autophagy, leading to desensitizing cells from protein aggregates, such as aggregates of mutant huntingtin-exon1. Thus, we propose an additive role of neddylation and HYPK in clearance of protein aggregates by autophagy, resulting in cytoprotective effect during proteotoxic stress.

## INTRODUCTION

Proteostasis is a dynamic process that ensures systemic maintenance of quality and quantity of cellular proteome (**1, 2**). Stress-induced proteins’ misfolding and aggregation are countered by protein quality control systems (PQCS) (**3**) which comprise a complex set of mechanisms, starting from guided folding and oligomerization of nascent polypeptides by chaperones (**4**) to ordered elimination of irreparable and non-functional proteins by proteasome and autolysosome machineries (**5, 6**). While the soluble proteins and smaller protein complexes are cleaved by 26S proteasomal system, the phase-separated insoluble protein aggregates are mainly degraded by autophagosomal/lysosomal pathway (**7**). Macroautophagy (hereafter referred as autophagy) is a general mechanism that devours cellular materials, such as damaged organelles, protein aggregates, intracellular pathogens etc., in a membranous vesicle, referred as autophagosome, followed by hydrolytic degradation of autophagosomal contents after fusion of autophagosome with lysosome (**8**). Initiation of autophagosome formation from phagophore is critically regulated by several protein complexes (MTOR complex, ULK1 complex, Vps34 complex, etc.) (**9**), whereas the maturation stage is more spontaneous with the help of LC3, Beclin-1 and different ATG proteins (**10**).

Destabilization of cellular proteostasis by intrinsic or extrinsic stress causes formation of heterogenous protein aggregates (**11**) which are selectively cleared by autophagy, often the process is termed as aggrephagy (**7**). Ubiquitination of proteins is the activation signal for degradation of protein aggregates by canonical autophagy pathway (**12**). The covalent linkage of ubiquitin to the target proteins results from sequential activation and functioning of E1, E2 and E3 ligases (**13**). Lysine-63 (K63) type polyubiquitin-linked protein aggregates are subjected to autophagic clearance (**14**). Although ubiquitination is the major and global modulator of aggrephagy, ubiquitin-like modifiers (UBLs) are also speculated to have contrasting functions in stress-induced autophagy. While the small ubiquitin-related modifier 1 (SUMO1) suppresses autophagy (**15**), it is not known if other UBLs have regulatory functions in autophagy. Particularly, understanding the function of Nedd8, which is a closely related protein to ubiquitin, in proteostasis is not well characterized. Conjugation of Nedd8 to its substrate proteins, mostly the cullin proteins (**16**), is mediated by its own set of ligase proteins (**17**). Interestingly, neurodegenerative disease-related proteins (stress granule proteins, inclusion body proteins etc.) are also neddylated (**18, 19**), indicating that neddylation of aggregation-prone proteins functions in clearance of protein aggregates. During proteotoxic stress, neddylation also sequesters the vulnerable nuclear proteins into aggregates (**20**). Moreover, thermal and redox stress also increase the cellular level of polyneddylated proteins (**21**). While such neddylated proteins undergo NUB1-assisted turnover by proteasomal system (**22**), the autophagy modulatory functions of Nedd8 remain unclear.

Autophagy receptor proteins are integral components in autophagosomal compartmentalization of protein aggregates (**23**). Other than binding to ubiquitin or UBLs, autophagy receptors also recruit LC3, leading to further assembly of autophagosomal machinery to phagophores. SQSTM1 (**24**), TAX1BP1 (**25**), TOLLIP (**26**) etc., are known ubiquitination-dependent autophagy receptors that promote aggrephagy. However, the autophagy receptors of other UBLs are not well defined. In this context, function of a member of intrinsically unstructured chaperone-like proteins (IUPs), named HYPK, in autophagy modulation appears promising. HYPK functions in sequestering and reducing aggregates of different cytosolic and nuclear proteins (**27, 28**). Previous studies from our group have showed that the balance of structural convolution of HYPK is maintained by complex intra- and inter-molecular interactions of its hydrophobic region, low complexity region and disordered nanostructure (**29, 30**). Earlier studies found a putative acetyltransferase aiding function of HYPK at ribosome (**31**). In *Caenorhabditis elegans*, HYPK forms proteasome blocking complexes (**32**), with a consequence in aging of the organism (**33**).

In this article, we report the regulatory functions of UBLs, specifically Nedd8, and HYPK in proteotoxic stress-induced aggrephagy. Assays for functional analysis reveal that polyneddylation of proteins serves as a post-translational modification (PTM) mark for clearance of protein aggregates by autophagy. In this pathway, HYPK functions as an autophagy receptor that scaffolds Nedd8 and LC3 by using its ubiquitin-associated (UBA) domain and LC3 interacting region (LIR). Both neddylation of proteins and HYPK act as aggrephagy inducer by helping autophagosome biogenesis around protein aggregates, such as those arising from defective ribosomal products of puromycin-treated or aggregation-prone huntingtin-exon1 expressing cells.

## RESULTS

### siRNA screen for ubiquitin-like proteins that modulate autophagic degradation of proteins during proteotoxic stress

The existing and newly synthesized proteins are vulnerable to proteotoxic stress (**34**). Chemical induction of intracellular proteotoxicity by puromycin occurs due to premature translation failure (**35**), resulting in formation of defective ribosomal products (**36**) that are primarily cleared from cells by autophagic degradation (**37**). While it is known that proteotoxic stress-induced protein aggregates are ubiquitinated prior to their autophagic degradation, the role of other ubiquitin-like proteins (UBLs) (**Supporting figure 1**) in this process is less understood. Hence, a screening of UBLs can shed more light on their functions in autophagy. In order to conduct the screen of UBLs that drive the autophagic degradation of proteins, we first determined the conditions of puromycin-induce proteotoxicity that transduced its effects by increasing the autophagy flux without causing cell death. Autophagy flux was monitored by a cell assay that measured the tandem fluorescence of RFP-GFP-LC3B (expressed in a stable cell-line of MCF7) in different conditions. This is a standard autophagy flux assay in which RFP-GFP-LC3B shows green (due to GFP) and red (due to RFP) double fluorescence in autophagosomes (AVs) and only red fluorescence in autolysosomes (ALs), indicating spatiotemporal progress of formation and maturation of AVs. Treatment of 15μg/ml puromycin (this concentration of puromycin was used in all experiments unless otherwise stated) to cells significantly increased the number of GFP+/RFP+ LC3B (AVs) and GFP-/RFP+ LC3B (ALs) puncta than untreated cells **(Figure 1A).** Conversion of native LC3B-I to LC3B-II also increased during puromycin treatment **(Figure 1B),** indicating high autophagy flux. Thus, we chose puromycin as a reproducible inducer of proteotoxic stress and autophagy in cells.

**Figure 1.**
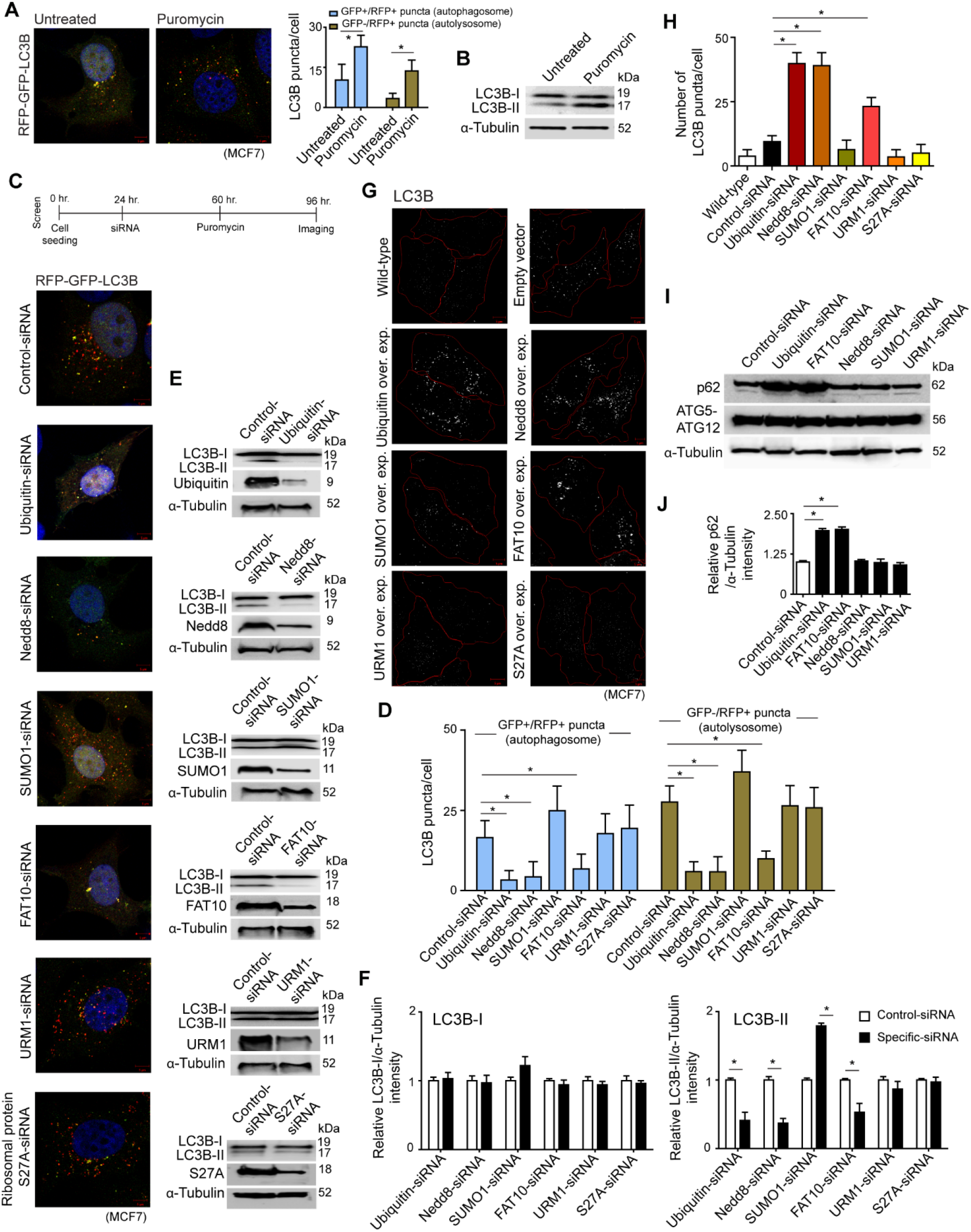
siRNA screen for autophagy modulatory ubiquitin-like proteins in proteotoxic stress. (A) Stable RFP-GFP-LC3B expressing MCF7 cells were untreated or treated with puromycin [15μg/ml] for 24 hours. Left: confocal fluorescence microscopy images of GFP+/RFP+ LC3B autophagosomes and GFP-/RFP+ LC3B autolysosomes. Right: Quantitative count [mean ± SD] of GFP+/RFP+ LC3B autophagosomes and GFP-/RFP+ LC3B autolysosomes; ~ 100 cells analysed in each sample [* P<0.05]. (B) Representative image of immunoblot of LC3B in lysate of untreated and puromycin-treated MCF7 cells. (C and D) Stable RFP-GFP-LC3B expressing MCF7 cells transfected with different ubiquitin-like protein-specific siRNAs and subsequent treatment of cells with puromycin for 36 hours. C: timeline of the screening experiment and confocal fluorescence microscopy images of GFP+/RFP+ LC3B autophagosomes and GFP-/RFP+ LC3B autolysosomes. D: mean ± SD number of GFP+/RFP+ LC3B autophagosomes and GFP-/RFP+ LC3B autolysosomes; ~ 150 cells analysed in each experiment [* P<0.05]. (E and F) E: representative immunoblot of LC3B from lysate of control-siRNA and UBL-specific siRNA-treated MCF7 cells. F: densitometric quantification of LC3B-I and LC3B-II bands relative to α-tubulin of blots [* P<0.05]. (G and H) MCF7 cells transfected with individual UBL-expressing clone. G: representative confocal immunofluorescence images of LC3B puncta. H: average ± SD number of LC3B puncta/cell in different UBL overexpression condition; ~ 150 cells analysed in each experiment [* P<0.05]. (I and J) Effect of knockdown of UBLs on expression level of autophagy receptor p62 and autophagic proteins ATG5, ATG12. I: representative immunoblots of p62 and ATG5-ATG12 in lysate of different UBL knockdown cells. J: densitometric quantification of p62 bands relative to α-tubulin in immunoblots. [α-tubulin is loading control in immunoblots. Scale bar in confocal microscopy images represents 5μm. All the presented microscopy and immunoblot data are representative of at least three independent experiments.]

We screened siRNA combinations (6-18 siRNA/gene, 10nM) against six UBLs, including the ubiquitin-siRNA (positive control), and a negative control-siRNA to understand the potency of the UBLs in modulating autophagy during proteotoxic stress (timeline of screening experiment is in **Figure 1C).** Other than the ubiquitin-siRNA, two siRNAs targeting the Nedd8 and FAT10 showed repression of autophagy flux. Nedd8-siRNA and FAT10-siRNA prevented the formation of GFP+/RFP+ AVs and GFP-/RFP+ ALs compared to control-siRNA during proteotoxic stress **(Figure 1C, 1D).** These two siRNAs also reduced the conversion of LC3B-I to LC3B-II in normal growth condition **(Figure 1E, 1F).** Contrary to Nedd8 and FAT10, downregulation of SUMO1 by siRNA increased formation of AVs/ALs and conversion of LC3B-I to LC3B-II. Thus, Nedd8 and FAT10 showed autophagy inducing properties like ubiquitin, whereas SUMO1 had opposite effect during proteotoxic stress.

Ubiquitin, Nedd8 and FAT10 appeared to be activators of basal autophagy than other UBLs. Higher expression of these three proteins significantly increased autophagy as quantified by the formation LC3B puncta in cells **(Figure 1G, 1H).** The effect of Nedd8 in increasing the basal autophagy was similar to ubiquitin but higher than FAT10.

Ubiquitin-dependent autophagy of proteins involves the receptor function of p62/SQSTM1 (**38**). Downregulation of ubiquitin expression caused reduction of autophagy flux and concomitant accumulation of p62 in cells **(Figure 1I, 1J).** While knockdown of FAT10 showed similar phenotype, Nedd8 downregulation did not show such effect on p62 levels in cells **(Figure 1I, 1J).** Thus, Nedd8 apparently modulated autophagy of proteins in p62 independent manner, possibly with the involvement of other scaffolding protein(s).

### Proteotoxicity-induced Nedd8 granules are cleared by autophagy

The preliminary finding from the knockdown (KD) screen that Nedd8 is a positive modulator of autophagy led us to examine the crosstalk between protein neddylation and their autophagic removal during proteotoxic stress. We observed formation of Nedd8 granules in puromycin-treated cells, but not in the untreated cells **(Figure 2A).** The Nedd8 granules were high molecular weight neddylated protein aggregates **(Figure 2B).** This observation was in-line with a previous study that also showed similar accumulation of neddylated protein granules during proteotoxic stress (**20**). Interestingly, we found that many of the puromycin-induced Nedd8 granules were also positive for LC3B **(Figure 2C, 2D).** Inhibition of protein neddylation by a chemical inhibitor, MLN4924 (concentration: 1μM in culture medium of all experiments unless otherwise stated), not only abolished the formation of Nedd8 granules in puromycin-treated cells, but it also depleted the LC3B foci which were otherwise seen at the Nedd8 granules in puromycin-treated cells **(Figure 2C).** Accordingly, cotreatment of cells with puromycin and MLN4924 decreased the basal level conversion of LC3B-I to LC3B-II compared to cells that were treated only with puromycin **(Figure 2E).** These observations indicated that the neddylated protein granules could be undergoing autophagic compartmentalization during proteotoxic stress.

**Figure 2.**
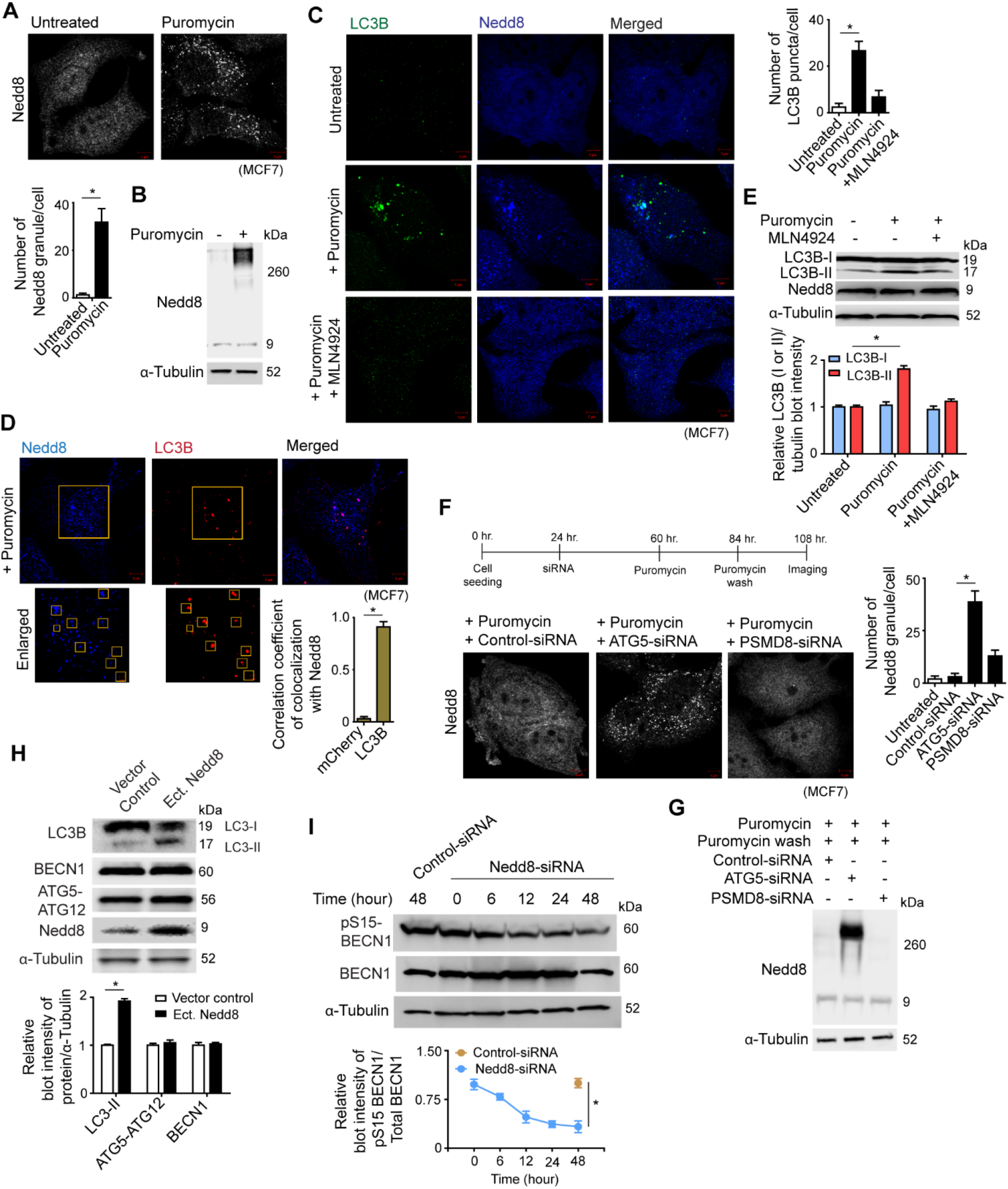
Nedd8 clears proteotoxic stress-induced protein aggregates by autophagy. (A and B) MCF7 cells untreated or treated with puromycin [15μg/ml] for 24 hours. A: confocal immunofluorescence microscopy images of Nedd8 granules, and mean ± SD quantification of Nedd8 granules; ~ 150 cells analysed in each sample [* P<0.05]. B: representative immunoblot of Nedd8 from cell extracts. (C) MCF7 cell were not treated or pretreated with 1μM MLN4924 for 12 hours and then exposed to puromycin [15μg/ml] for 24 hours. Confocal immunofluorescence microscopy images of Nedd8 and LC3B. Quantification of LC3B puncta with mean ± SD in cells; ~ 150 cells analysed in each sample [* P<0.05]. (D) Representative confocal immunofluorescence microscopy image of colocalization of Nedd8 granules and LC3B in puromycin-treated [15μg/ml] MCF7 cells. Inset depicts enlarged images of Nedd8 and LC3B positive puncta marked with yellow rectangles. Correlation coefficient of colocalization of Nedd8 granules with LC3B as compared to mCherry in MCF7 cells, ~ 50 cells analysed in each sample [* P<0.05]. (E) Representative immunoblot of LC3B and Nedd8 in the condition described as in (c), with densitometric quantification of the LC3B-I and LC3B-II bands relative to α-tubulin in immunoblots [* P<0.05]. (F and G) ATG5, PSMD8 and control knockdown MCF7 cells were treated with puromycin [15μg/ml], followed by washout of puromycin with normal cell culture medium as presented in timeline of the experiment. F: confocal immunofluorescence microscopy images of Nedd8, and mean ± SD quantification of Nedd8 granules in different knockdown cells, ~ 150 cells analysed in each sample [* P<0.05]. G: Representative immunoblot of Nedd8 from the cell lysate of control, ATG5 and PSMD8 knockdown MCF7 cells. (H) Representative immunoblot of autophagy marker proteins from lysate of control and Nedd8 overexpressing MCF7 cells. Densitometric quantification of bands of autophagy marker proteins relative to α-tubulin of blots [* P<0.05]. (I) Representative immunoblots of phosphoserine15-beclin-1 and beclin-1 [BECN1] in a time-chase experiment from lysate of control and Nedd8 knockdown MCF7 cells. Densitometric quantification of phosphoserine15-beclin-1 relative to beclin-1 in the immunoblots [* P<0.05]. [α-tubulin is loading control in immunoblots. Scale bars in confocal microscopy images represent 5μm and 1μm for enlarged images. All the presented microscopy and immunoblot data are representative of at least three independent experiments.]

Next, we examined if the neddylated granules were cleared through autophagy pathway by knocking down one or more proteins of core autophagy machinery. Although puromycin initiated the accumulation of Nedd8 granules, they were effectively cleared in control knockdown cells after the washout of puromycin **(Figure 2F, 2G).** On the contrary, a significant number of Nedd8 granules persisted in the ATG5 knockdown cells after puromycin washout **(Figure 2F, 2G).** Blocking the 26S proteasomal pathway by knockdown of an essential proteasomal protein, PSMD8, also showed accumulation of Nedd8 granules, even though almost five times lesser than the effect of ATG5 knockdown **(Figure 2F, 2G).** Taken together, these data suggest that proteotoxic stress causes formation of neddylated substrate granules that are cleared by autophagy pathway.

To confirm the role of Nedd8 in autophagy, we further studied the expression of autophagy markers during differential expression of Nedd8 protein. Higher expression of Nedd8 in MCF7 cells caused increases lipidation of LC3B-I to LC3B-II, signifying the activation of autophagy pathway **(Figure 2H).** Knockdown of Nedd8 temporally reduced the cellular level of phosphoserine15-BECN1 **(Figure 2I).** Such results provide the evidence that Nedd8 augments a noncanonical autophagy pathway to degrade proteins.

### Polyneddylated aggregation-prone huntingtin exon1 is degraded by autophagy

Having found that proteotoxicity-induced Nedd8 granules are cleared by autophagy, we were interested to find if polyneddylated proteins are in general subjected to autophagic degradation. Huntingtin is one of the few number of substrate proteins that undergo polyneddylation (**39**). Consistent with the previous studies, we observed that the polyglutamine-expanded mutant huntingtin exon1 (htt97Qexon1) was polyneddylated **(Figure 3A).** Denaturing (with 1% SDS) immunoprecipitation of Nedd8 (to purify only neddylated proteins and prevent purification of proteins that are noncovalently associated with Nedd8) followed by immunoblotting against htt97Qexon1 showed the high molecular weight neddylated htt97Qexon1 **(Figure 3A).** Neddylation of huntingtin exon1 aggregates in IMR-32 cells was also evident from the observation of high Nedd8 colocalization/deposition with htt97Qexon1 aggregates **(Figure 3B).** Since a nonspecific protein (blue fluorescent protein, BFP) did not colocalize with htt97Qexon1 **(Figure 3B),** it was apparent that neddylation of htt97Qexon1 was specific, and not just trapping of Nedd8 with the sticky huntingtin exon1 aggregates.

**Figure 3.**
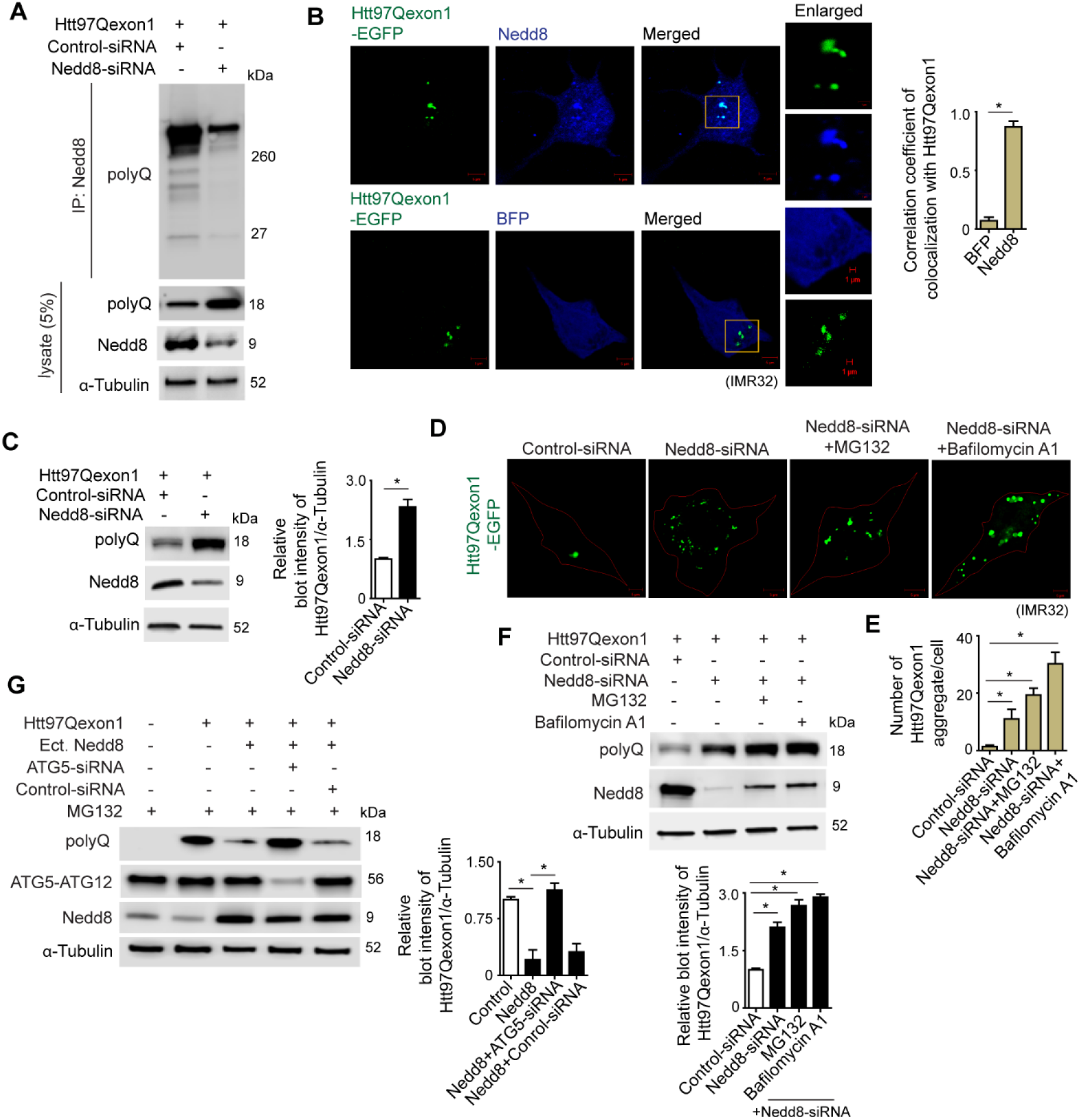
Neddylation mediates aggrephagy of mutant huntingtin exon1 protein aggregates. (A) Representative immunoblots of htt97Qexon [polyQ] and Nedd8 of denaturing immunoprecipitations of neddylated proteins from extract of stable htt97Qexon1 expressing IMR-32 cells transfected with control or Nedd8 siRNAs. (B) Confocal immunofluorescence image of colocalization of Nedd8 with htt97Qexon1-GFP in IMR-32 cells stably expressing htt97Qexon1-GFP. Quantification of colocalization coefficient of htt97Qexon1 GFP with Nedd8 compared to BFP, ~ 200 cells analysed in each sample [* P<0.05]. (C) Representative immunoblot of htt97Qexon1 and Need8 from lysate of htt97Qexon1-expressing IMR-32 cells transfected with control or Nedd8 siRNAs. Densitometric quantification of htt97Qexon1 bands relative to α-tubulin of blots [* P<0.05]. (D and E) Control or Nedd8 siRNA was transfected in stable htt97Qexon1-GFP expressing IMR-32 cells. Nedd8 knockdown cells were exposed to 5μM MG132 or 1μM bafilomycin-A1 for 24 hours. D: confocal fluorescence microscopy images of htt97exon1-GFP. E: quantification [mean ± SD] of htt97exon1-GFP aggregates, ~ 200 cells analysed in each sample [* P<0.05]. (F) Representative immunoblot of htt97Qexon1 and Nedd8 from experiments following the same procedure as that in (d), except that the IMR-32 cells stably expressed htt97Qexon1. Densitometric quantification of htt97Qexon1 bands relative to α-tubulin of blots [* P<0.05]. (G) Control or ATG5 siRNA was transfected in htt97Qexon1 and Nedd8 overexpressing IMR-32 cells in presence of 5μM MG132. Representative immunoblots of htt97Qexon1 and Need8. Densitometric quantification of htt97Qexon1 bands relative to α-tubulin of blots [* P<0.05]. [α-tubulin is loading control in immunoblots. Scale bars in confocal microscopy images represent 5μm. All the presented microscopy and immunoblot data are representative of at least three independent experiments.]

In theory, huntingtin exon1 could be subjected to three different types of neddylation - mononeddylation, polyneddylation and multi-mononeddylation. To understand the neddylation pattern of htt97Qexon1, we generated a Nedd8 mutant construct in which all the lysine residues of Nedd8 were mutated to arginine (Nedd8-allR) **(Supporting figure 2A).** This Nedd8-allR mutant cannot covalently attach to another Nedd8 or Nedd8-allR molecule through lysine-linkage, thereby preventing the formation of polyneddylated chains of Nedd8-allR. However, conjugation of Nedd8-allR to proteins can form mononeddylated and multi-mononeddylated substrates. Coexpression of htt97Qexon1 with Nedd8 or Nedd8-allR showed that high molecular weight conjugated complexes of htt97Qexon1 formed only in presence of Nedd8 **(Supporting figure 2B).** Nedd8-allR conjugation resulted in mononeddylated htt97Qexon1, but not the polyneddylated or multi-mononeddylated htt97Qexon1.

During stress or higher expression of Nedd8, polyubiquitin chains also incorporate Nedd8 (**40**). This phenomenon does not depend upon the Nedd8-activating enzyme (NAE1). Instead, Nedd8 is activated by the ubiquitin-activating enzyme (UBA1) (**41**). In our experiments, the homogeneity of polyneddylation of htt97Qexon1 was analyzed by using NAE1 and UBA1 inhibitors. Treatment of the htt97Qexon1 expressing IMR-32 cells with NAE1 inhibitor, MLN4924, decreased polyneddylation of the protein **(Supporting figure 2C).** Application of UBA1 inhibitor, MLN7243 (concentration: 1μM in culture medium of all experiments unless otherwise stated), to the cells had no effects on the polyneddylation of huntingtin exon1 **(Supporting figure 2C).**

To understand the effect of neddylation on huntingtin exon1 degradation by autophagy, we measured the cellular level of htt97Qexon1 in varying expression conditions of Nedd8. Knockdown of Nedd8 effectively decreased the degradation of htt97Qexon1 compared to control cells **(Figure 3C).** Number of htt97Qexon1 aggregates also increased in the Nedd8-KD cells **(Figure 3D, 3E).** Exposure of htt97Qexon1 expressing cells (in Nedd8-KD background) with proteasomal inhibitor (MG132; concentration: 5μM in culture medium of all experiments unless otherwise stated) or autophagy inhibitor (Bafilomycin-A1; concentration: 1μM in culture medium of all experiments unless otherwise stated) showed further increase of htt97Qexon1 protein and its aggregates in terms of their numbers and size **(Figure 3D, 3E, 3F).** In this respect, blocking the autophagy pathway had a more pronounced effect than blocking the proteasomal pathway. Previous reports found that neddylated huntingtin could be cleared through proteasomal pathway by the activity of NUB1 protein (**39**), possibly through the activity of p97^UFD1/NPL4^ Complex (**42**). However, given the condition that polyubiquitinated proteins, such as huntingtin, could be redundantly degraded by both proteasomal and autophagosomal pathways (**43**), we tested if polyneddylated htt97Qexon1 was also subjected to autophagosomal degradation. While higher expression of Nedd8 enhanced the clearance of htt97Qexon1 from cells, downregulation of ATG5 prevented such Nedd8-facilitated clearance of htt97Qexon1 **(Figure 3G).** MG132 treatment to cells ensured that neddylation-dependent degradation of htt97Qexon1 was not occurring through proteasomal pathway. Since Nedd8 could assist the degradation of htt97Qexon1 in presence of MG132, it was reasonable to understand that Nedd8 had regulatory functions in the autophagic degradation of mutant huntingtin exon1. Overall, these observations showed that Nedd8 is an autophagy modulating protein, and polyneddylation can serve as a signal of autophagic degradation of protein aggregates.

### HYPK is the receptor in neddylation-dependent autophagy

The canonical degradation of polyubiquitinated proteins by autophagy requires the scaffolding function of p62/SQSTM1 protein. p62 has a UBA and LC3 interacting region (LIR) which are used by the protein to interact with ubiquitin and LC3 respectively. We expected that a protein with similar scaffolding function would be required in the autophagy of polyneddylated proteins. The finding that cellular level of p62 was unchanged during inhibition of neddylation-dependent autophagy (described in first section of the results; **Figure 1I, 1J),** led us to turn our attention to identify the scaffolding protein which is involved in the interaction with Nedd8 and LC3.

#### (I) HYPK interacts with Nedd8 through its C-terminal UBA domain

Based on the fact that UBA domains recognize ubiquitin and UBLs (**44**), we tested a knockdown screen for the effect of thirty UBA domain-containing proteins in retention of cellular Nedd8 granules. Downregulation of five UBA domain-containing proteins significantly increased the number of polyneddylated substrate granules in cells **(Figure 4A, 4B);** with HYPK knockdown showing highest accumulation of neddylated granules **(Figure 4A).** To get further idea of an ideal binding of Nedd8 with the UBA domains, we conducted computational docking/binding studies of Nedd8 with different UBA domains of selected proteins. A higher free energy of assembly dissociation [ΔG(diss)] would indicate stronger binding of Nedd8 with the respective UBA domain. The binding affinity of one of the UBA domains of NUB1 with Nedd8 was highest, followed by the binding of HYPK-UBA with Nedd8 **(Figure 4C),** implying that UBA domains of NUB1 and HYPK form thermodynamically stable complexes with Nedd8.

**Figure 4.**
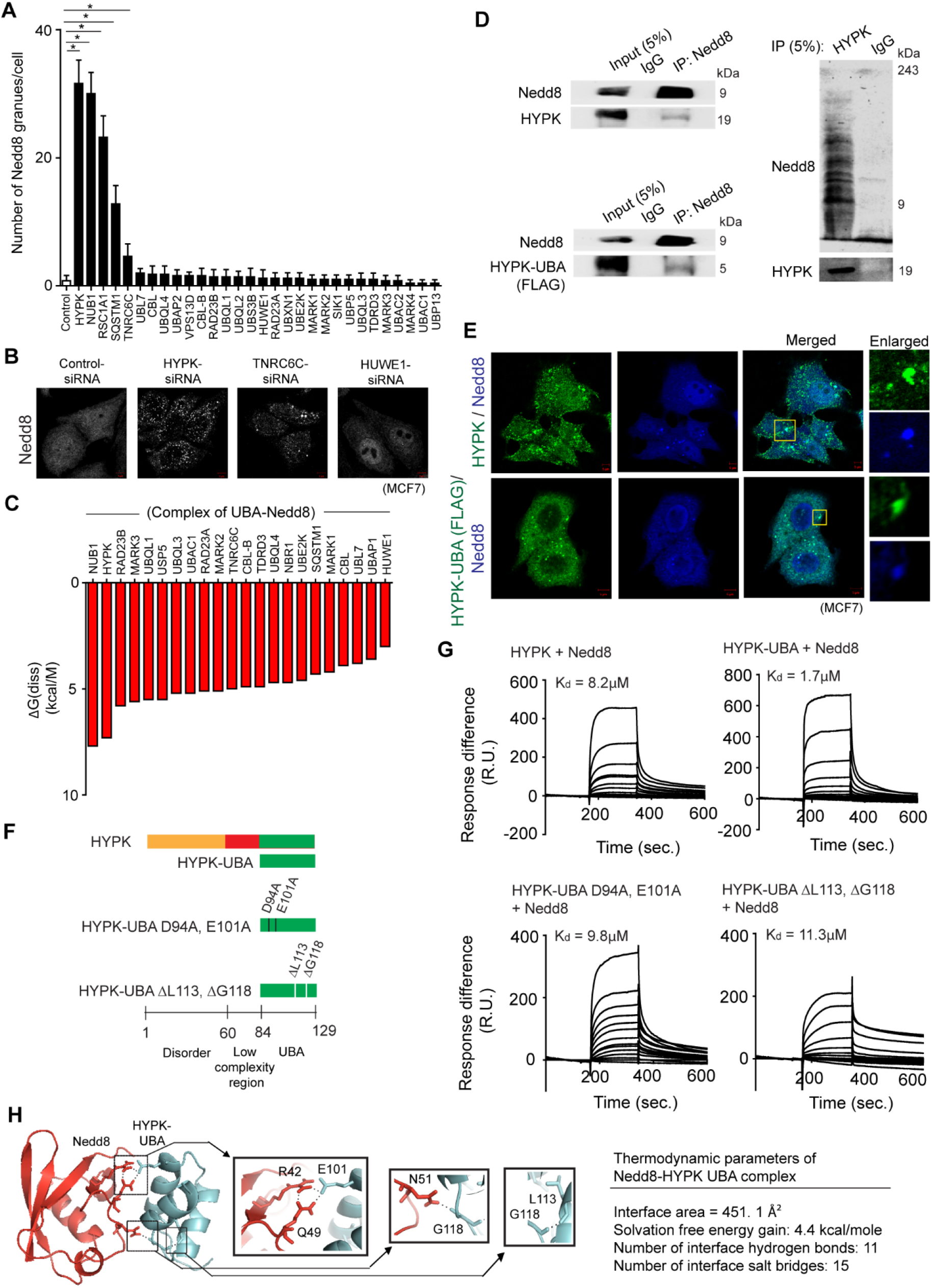
HYPK binds to Nedd8 by its UBA domain. (A and B) MCF7 cells were transfected with control or siRNA against different UBA domain containing proteins. A: number [mean ± SD] of Nedd8 granules in cells, ~ 300 cells analysed in each sample [* P<0.05]. B: Representative confocal immunofluorescence microscopy images of Nedd8. (C) *In silico* molecular docking of Nedd8 was done on UBA domains of selected human proteins. Free energy of dissociation of assemblies/complexes was computed in PISA webserver. (D) Representative immunoblots in non-denaturing conditions of – Left: Nedd8, HYPK and FLAG-tagged HYPK-UBA of immunoprecipitation of Nedd8 from cell lysate of MCF7 cells [FLAG-tagged HYPK-UBA was ectopically expressed]; Right: HYPK and Nedd8 of immunoprecipitation of HYPK from cell lysate of MCF7 cells. (E) Confocal immunofluorescence microscopy images of Nedd8, HYPK and FLAG-tagged HYPK-UBA in MCF7 cells. (F) Representation of HYPK, its UBA domain and different mutants of HYPK-UBA domain that were used in this section of study. HYPK-UBA domain contained the C-terminal 45 amino acid region; the D94 and E101 residues were mutated to alanine in HYPK-UBA D94A, E101A construct; the L113 and G118 residues were deleted in HYPK-UBA ΔL113, ΔG118 construct. (G) Quantitative binding responses and affinities of interactions between recombinant HYPK/HYPK-UBA/ HYPK-UBA D94A, E101A/HYPK-UBA ΔL113, ΔG118 and Nedd8 probed by surface plasmon resonance assays. Dissociation constants [K_d_ value] were calculated by using 1:1 Langmuir model of binding. (H) Structural basis and thermodynamic parameters of interactions between Nedd8 and HYPK-UBA in the HYPK-Nedd8 complex. Left: predicted structure of complex of HYPK and Nedd8. Right: the conserved residues in HYPK-UBA and their interaction partners in Nedd8 are shown with stick representation. Hydrogen bonds are highlighted with black dotted line. [α-tubulin is loading control in immunoblots. Scale bars in confocal microscopy images represent 5μm. All the presented microscopy and immunoblot data are representative of at least three independent experiments.]

The functional binding of the UBA domain of NUB1 to Nedd8 (**45**) is known to be involved in NUB1-facilitated degradation of neddylated proteins through proteasomal pathway. On the other hand, aggregate-sequestering function of HYPK is linked to the degradation of the aggregation-prone proteins in an unknown pathway (**27**). Thus, it is crucial to understand how HYPK delivers the protein aggregates to degradation system.

HYPK structure shows the existence of a putative UBA domain in its C-terminal region (**30**). This region is well conserved in the HYPK proteins of different organisms **(Supporting figure 3A).** A multiple sequence alignment-based phylogenetic analysis to cluster the similar UBA domains of human proteins directed to the observation that the sequence of the second UBA domain (UBA2) of NUB1 protein is closest to the sequence of HYPK-UBA domain **(Supporting figure 3B).** Thus, we tested if HYPK is a Nedd8 interacting protein. HYPK could strongly bind to Nedd8 by using its UBA domain *in vitro* conditions. Immunoprecipitation of Nedd8 from MCF7 cell lysate in non-denaturing condition, followed by immunoblotting showed that both HYPK and HYPK-UBA could be pulled-down with Nedd8 **(Figure 4D).** A reciprocal immunoprecipitation of HYPK, followed by immunoblotting for Nedd8 and HYPK was also done to test which form of Nedd8 binds to HYPK. The immunoblot profile showed HYPK interaction with both monomeric Nedd8 and polyneddylated chain **(Figure 4D).** Furthermore, we also observed that HYPK and HYPK-UBA signals disappeared in the blots of denaturing (with 1% SDS) immunoprecipitation of Nedd8 **(Supporting figure 3C),** indicating that HYPK and HYPK-UBA were not neddylated, but they were noncovalently bound to Nedd8. HYPK and HYPK-UBA were also observed to show high spatial colocalization with intracellular Nedd8 **(Figure 4E).**

We identified a set of critical residues in HYPK-UBA domain [aspartate-94 (D94), glutamate-101 (E101), leucine-113 (L113) and glycine-118 (G118); residue positions were numbered according to human HYPK isoform-1; NCBI accession number: NP_057484.3] that were required for strong and efficient binding of HYPK to Nedd8. These residues are typically conserved in the HYPK proteins of different organisms **(Supporting figure 3A).** In order to understand the function of these amino acids in HYPK binding to Nedd8, we generated different mutants of HYPK-UBA in which the conserved residues were either mutated or deleted **(Figure 4F).** In the HYPK-UBA D94A, E101A construct, the D94 and E101 residues were mutated to alanine; whereas L113 and G118 residues were deleted in the HYPK-UBA ΔL113, ΔG118 mutant. In the protein-protein interaction assays by surface plasmon resonance (SPR), both the mutants displayed lower binding affinity for Nedd8 compared to the affinity of wild-type HYPK and HYPK-UBA for Nedd8 **(Figure 4G).** It implied that the four conserved residues in HYPK-UBA were necessary, but not sufficient, for HYPK-UBA interaction with Nedd8. Although the conserved amino acids in HYPK-UBA were critical for HYPK binding to Nedd8, the neighboring residues possibly provided additional support to the interaction. We also noted that the full-length HYPK protein had lower binding affinity for Nedd8 than HYPK-UBA. The reduced binding affinity of HYPK for Nedd8 could be due to the intramolecular interaction between the disorder nanostructure and low complexity region (LCR) of HYPK, as shown in one of our previous studies (**29**). Such an intramolecular interaction can interfere in the optimal binding of HYPK-UBA with Nedd8.

In the available crystal structure of HYPK (**46**), the E101 and G118 residues are exposed on the surface of the protein, allowing these two residues to directly interact with the proximal residues of Nedd8 in the HYPK-Nedd8 complex **(Figure 4H).** On the other hand, D94 and L113 residues are located at the interface of two helices of the globular core of HYPK-UBA domain **(Figure 4H).** Thus, these residues are more likely to be involved in stabilization of the HYPK-UBA domain rather than making direct interactions with Nedd8.

#### (II) HYPK is an LC3 interacting protein

To understand if HYPK is involved in facilitating the autophagic degradation of neddylated proteins, we investigated the interaction of HYPK with LC3. HYPK lacks any conventional LIR which could be represented by [W/Y/F]xx[L/V/I] sequence. However, an unbiased analysis of HYPK sequence predicted a putative atypical LIR sequence represented by Y_49_AEE_52_ **(Figure 5A).** Such an atypical LIR with an aromatic amino acid at first position and an acidic amino acid (instead of a hydrophobic amino acid) at the fourth position is previously reported in the UBA5 protein (in the UBA5, the LIR is WGIE) (**47**). The tyrosine-49 (Y49) is also conserved in the HYPK protein of various organisms **(Figure 5A)**

**Figure 5.**
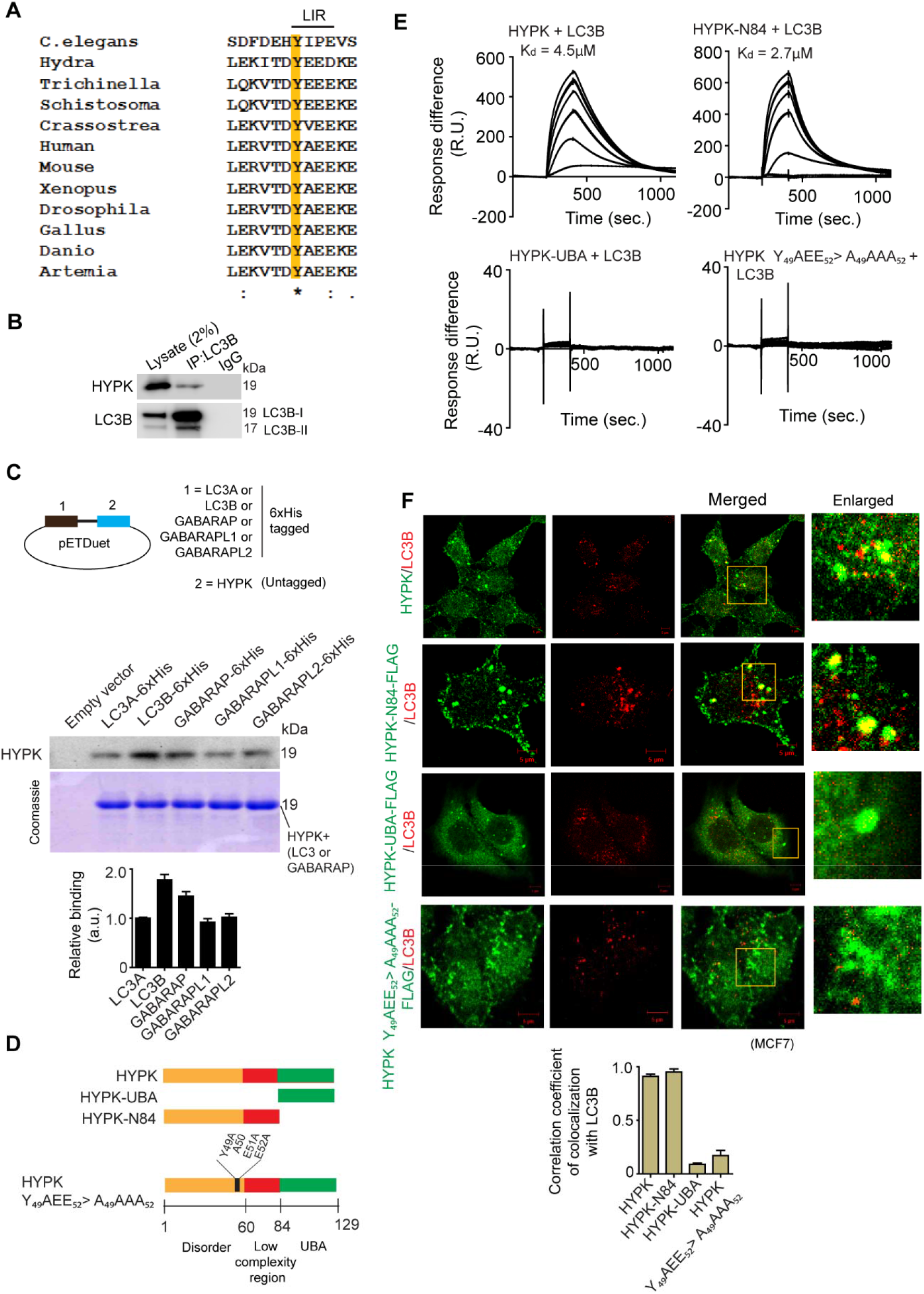
HYPK is a LC3 binding protein. (A) Multiple sequence alignment of the putative LC3 interacting region sequences of HYPK of different organisms. (B) Representative immunoblots of HYPK and LC3B from the non-denaturing immunocomplexes of endogenous LC3B from cell lysate of MCF7 cells. (C) Top: schematic representation of cloning of LC3A/LC3B/GABARAP/GABARAPL1/GABARAPL2 and HYPK in pETDuet-1 vector. Bottom: protein-protein interaction analysis by Ni-NTA binding of recombinant 6xhistidin-tagged LC3A/LC3B/GABARAP/ GABARAPL1/GABARAPL2 and untagged HYPK coexpressed in BL21DE3 strain of *Escherichia coli* from pETDuet-1 vector. Immunoblot of HYPK and coomassie blue staining of interacting pairs; empty vector was used for negative control. (D) Representation of HYPK and its mutants used in this part of study. HYPK-UBA was the C-terminal 45 residue region, HYPK-N84 was the N-terminal 84 amino acid region, the 49^th^, 51^st^ and 52^nd^ residues of HYPK were mutated to alanine in HYPK Y_49_AEE_52_>A_49_AAA_52_. (E) Quantitative analysis of binding interaction and affinities between recombinant HYPK/HYPK-UBA/HYPK-N84/HYPK Y_49_AEE_52_>A_49_AAA_52_ and LC3B determined by surface plasmon resonance assays. Dissociation constants [K_d_ value] were calculated by using 1:1 Langmuir model of binding. (F) MCF7 cells were transfected with HYPK/HYPK-UBA/HYPK-N84/HYPK Y_49_AEE_52_>A_49_AAA_52_. Confocal immunofluorescence microscopy images of HYPK, FLAG-tagged HYPK-UBA, FLAG-tagged HYPK-N84 and FLAG-tagged HYPK Y_49_AEE_52_>A_49_AAA_52_ and LC3B. Quantitative estimation of colocalization coefficient between proteins; ~ 200 cells analysed in each sample. [α-tubulin is loading control in immunoblots. Scale bars in confocal microscopy images represent 5μm. All the presented microscopy and immunoblot data are representative of at least three independent experiments.]

A non-denaturing immunoprecipitation assay with endogenous LC3B showed that HYPK was coimmunoprecipitated with LC3B **(Figure 5B).** This observation was comparable with the findings of another recent study that also reported proximity-dependent HYPK interaction with ATG8/LC3 in plants (**48**). To validate if HYPK directly interacts with LC3 (LC3A and LC3B) and the GABARAP class of proteins (GABARAP, GABARAPL1 and GABARAPL2), we conducted protein-protein interaction studies by pull-down assays using recombinant proteins. We coexpressed 6xhistidine-tagged LC3A/LC3B/GABARAP/GABARAPL1/GABARAPL2 and untagged HYPK in BL21DE3 strain of *Escherichia coli* by using pETDuet-1 vector (pETDuet-1 vector allows coexpression of two recombinant proteins in T7 promoter expression system). Nickel-NTA beads affinity pull-down showed copurification of HYPK with all the LC3 and GABARAP proteins **(Figure 5C),** suggesting that HYPK could globally interact with LC3 and GABARAP subfamily proteins.

To probe the function of putative LIR of HYPK in LC3 binding, we used truncation mutations and alanine scanning of HYPK as described in **Figure 5D.** The HYPK-UBA contained the C-terminal UBA domain and HYPK-N84 construct was comprised of the N-terminal eighty-four amino acids (including the putative LIR). The HYPK Y^49^AEE^52^>A^49^AAA^52^ was a site-directed mutagenesis construct in which all the four residues in 49^th^-52^nd^ amino acid region (YAEE) were mutated to alanine (AAAA) in the full-length HYPK. The binding assays of recombinant proteins by SPR showed that HYPK and HYPK-N84, but not the HYPK-UBA and HYPK Y^49^AEE^52^>A^49^AAA^52^, could bind to LC3B **(Figure 5E),** confirming that the LC3 binding site of HYPK is located in the N-terminal region which contains the LIR at 49^th^-52^nd^ amino acid stretch. To validate the intracellular association of HYPK with LC3B, we traced the colocalization of HYPK, HYPK-N84, HYPK-UBA and HYPK Y^49^AEE^52^>A^49^AAA^52^ with LC3B protein in MCF7 cells. Similar to the observations of SPR study, only HYPK and HYPK-N84 showed high colocalization with LC3B **(Figure 4F).** These results clearly show that HYPK interacts with LC3 by its atypical tyrosine-type (Y-type) LIR.

### HYPK modulates autophagy

Having found that HYPK could interact with LC3 and Nedd8, we explored if HYPK modulated the intracellular induction and flux of autophagy. Knockdown of HYPK reduced the basal level of cellular autophagy, as seen by reduction of LC3B-II in HYPK-siRNA-treated cells compared to control cells **(Figure 6A).** HYPK knockdown also reduced the number of LC3B puncta, whereas HYPK overexpression increased the number of LC3B puncta in MCF7 cells **(Figure 6B),** indicating that HYPK helps in formation (i.e. induction) of autophagosomes. Imaging of HYPK overexpressing cells by transmission electron microscopy also showed increased autophagosome formation **(Figure 6C),** while the control cells contained significantly lesser number of autophagosomes **(Figure 6C).**

**Figure 6.**
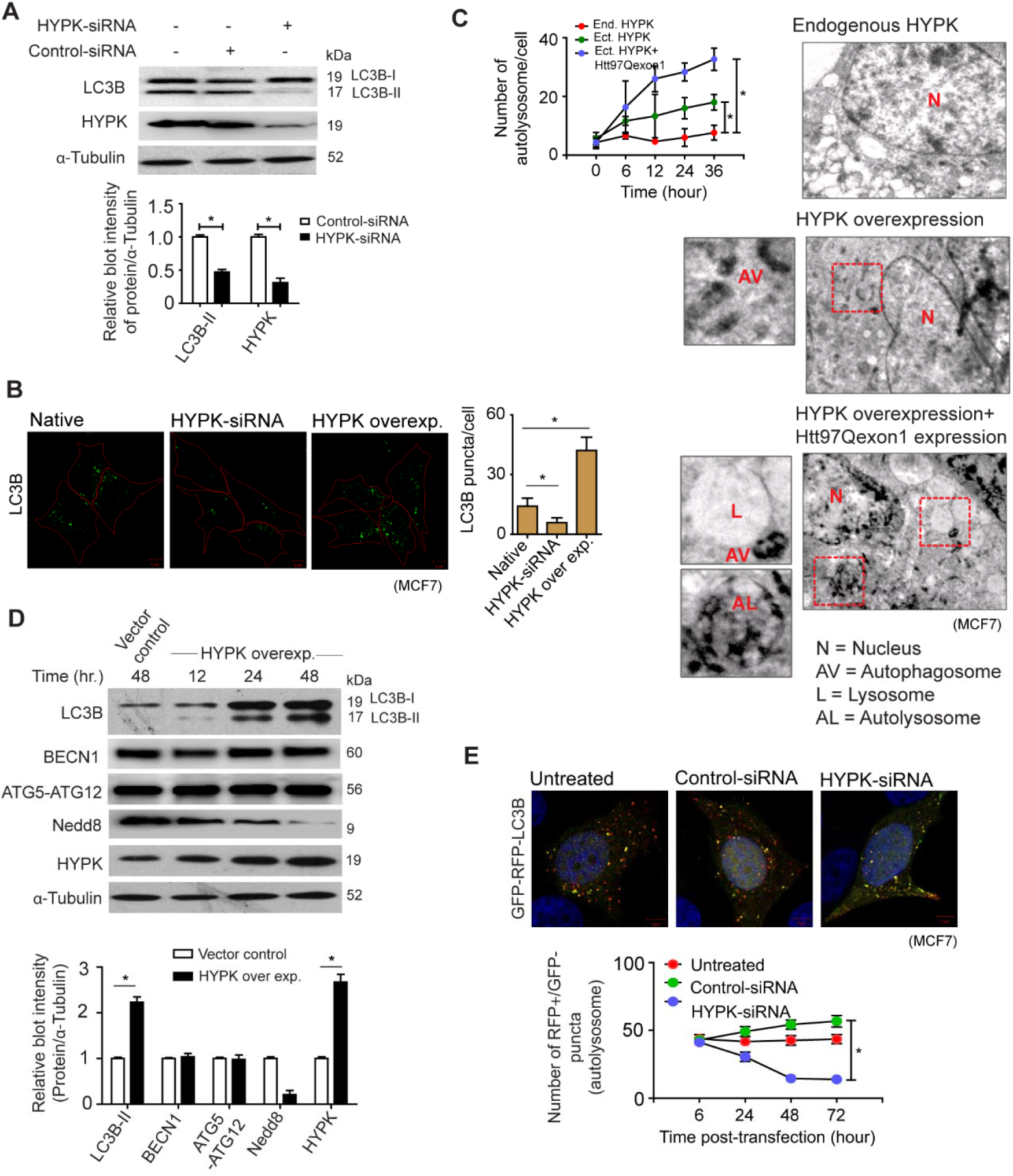
HYPK regulates initiation and flux of basal autophagy. (A) MCF7 cells were untransfected or transfected with control or HYPK siRNAs. Representative immunoblot of LC3B and HYPK from cell lysate. Densitometric quantification of LC3B-II and HYPK bands relative to α-tubulin of blots [* P<0.05]. (B) MCF7 cells were untransfected or transfected with HYPK siRNA or HYPK overexpressing clone. Confocal immunofluorescence microscopy images of LC3B puncta representing potential autophagosomes in cells. Quantification [mean ± SD] of number of LC3B puncta in cells; ~ 200 cells analysed in each sample [* P<0.05]. (C) Representative transmission electron micrographs of ultra-structures in MCF7 cells. Cells in the -upper panel: untreated, middle panel: transfected with HYPK overexpressing clone, lower panel: transfected with HYPK and htt97Qexon1 overexpressing clones. Quantitative [mean ± SD] estimation of autolysosomes in the above-mentioned cells for a timescale as mentioned in the figure, ~ 150 cells analysed in each sample [* P<0.05]. (D) MCF7 cells were transfected with empty vector or HYPK overexpressing clones. Representative immunoblot of LC3B, BECN1, ATG5, NEdd8 and HYPK of time-chase experiments from cell lysate. Densitometric quantification of bands of above-mentioned proteins relative to α-tubulin of blots [* P<0.05]. (E) Stable RFP-GFP-LC3B expressing MCF7 cells were untransfected or transfected with control or HYPK siRNAs. Confocal fluorescence microscopy images of GFP+/RFP+ LC3B autophagosomes and GFP-/RFP+ LC3B autolysosomes. Quantitative count [mean ± SD] of GFP-/RFP+ LC3B autolysosomes; ~ 100 cells analysed in each sample [* P<0.05]. [α-tubulin is loading control in immunoblots. Scale bars in confocal microscopy images represent 5μm. All the presented microscopy and immunoblot data are representative of at least three independent experiments.]

To further understand if HYPK was also involved in maintaining the autophagy flux, we conducted a time-chase experiment to monitor the cellular levels of LC3B-II, BECN1 and ATG5-ATG12 during overexpression of HYPK. HYPK maintained the steady-state flux of autophagy by continuously stimulating the lipidation of LC3B-I to LC3B-II without changing the cellular level of BECN1 and ATG5-ATG12 **(Figure 6D).** Additionally, the maturation of autophagosomes to autolysosomes decreased in HYPK knockdown cells. The count of RFP+/GFP-LC3B puncta (ALs) were almost four-fold less in HYPK knockdown cells than control cells **(Figure 6E).** Thus, with these results, it was evident that HYPK had enhancing function in selective autophagy induction and maturation.

### HYPK facilitates neddylation-dependent aggrephagy during proteotoxic stress

To determine if HYPK clears the neddylated protein aggregates by autophagy during proteotoxic stress, we analyzed the pattern of cellular distribution of HYPK in puromycin-treated cells. We observed a time-dependent increase of HYPK foci, which colocalized with Nedd8 granules, in puromycin-treated MCF7 cells **(Figure 7A, 7B).** It is, in fact, that we had previously shown the self-oligomerization of HYPK upon heterogenous proteins aggregates (such as aggregates of α-Synuclein-A53T, SOD1-G93A, htt97Qexon1 etc.) to sequester those protein aggregates (**27**). Complementary results in study showed that the formation of HYPK foci is dependent upon neddylation-status of proteins in puromycin-treated cells. HYPK could not form oligomeric granules in Nedd8-KD cells relative to control cells **(Figure 7C, 7D).** Again, this result was consistent with the previously observed phenomenon that the metastable states of UBA domain was required for self-association of HYPK (**30**).

**Figure 7.**
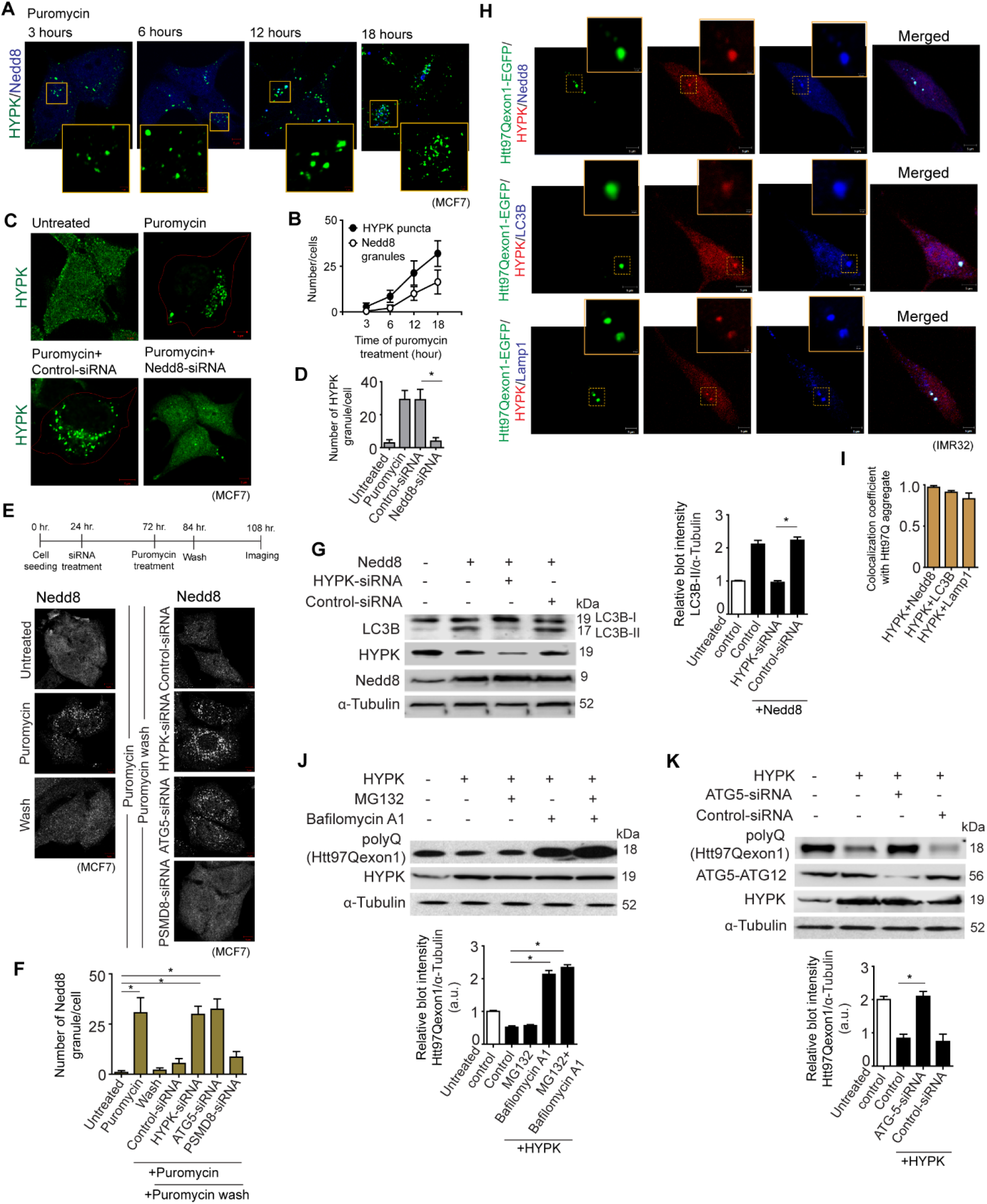
HYPK clears toxic aggregates of htt97Qexon1 by polyneddylation-dependent aggrephagy. (A) MCF7 cells were treated with puromycin [15μg/ml] for 0-18 hours. Confocal immunofluorescence microscopy images of HYPK and Nedd8. Inset shows enlarged region of HYPK and Nedd8 colocalized granules marked with yellow rectangles. (B) Quantification of HYPK and Nedd8 positive granules in the MCF7 cells with conditions as described in (a), ~ 100 cells analysed in each sample. (C) MCF7 cells were untreated or treated with [15μg/ml] puromycin for 24 hours. Puromycin treatment was done in untransfected or control or Nedd8 siRNA-transfected cells. Confocal immunofluorescence microscopy images of HYPK. (D) Quantification [mean ± SD] of HYPK granules in MCF7 cells with conditions as described in (c), ~ 100 cells analysed in each sample [* P<0.05]. (E and F) MCF7 cells were untransfected or transfected with control or HYPK or ATG5 or PSMD8 siRNA. Cells were treated with [15μg/ml] puromycin, followed by washout of puromycin as described in the timeline of the experiment. E: confocal immunofluorescence microscopy images of Nedd8. F: quantification of Nedd8 granules in cells with conditions as described in (e), ~ 100 cells analysed in each sample [* P<0.05]. (G) Control or HYPK siRNA was transfected in Nedd8 overexpressing MCF7 cells. Left: representative immunoblots of LC3B, HYPK and Nedd8 from the lysates of untransfected or transfected cells. Right: Densitometric quantification of LC3B-II bands relative to α-tubulin of blots [* P<0.05]. (H) Confocal immunofluorescence microscopy images of HYPK, Nedd8, LC3B and Lamp1 in stable htt97Qexon1-GFP expressing IMR-32 cells. (I) Coefficient of colocalization of HYPK, Nedd8, LC3B and Lamp1 with htt97Qexon1-GFP, ~ 100 cells analysed in each sample. (J) 5μM MG132 or 1μM bafilomycin-A1 was separately given for 24 hours to stable htt97Qexon1 expressing IMR-32 cells that were untransfected or transfected with HYPK overexpressing clone. Representative immunoblots of htt97Qexon1 and HYPK. Densitometric quantification of htt97Qexon1 bands relative to α-tubulin of blots [* P<0.05]. (K) Control or ATG5 siRNA were transfected in stable htt97Qexon1 expressing IMR-32 cells that also had HYPK overexpressing clone. Representative immunoblot of htt97Qexon1, ATG5-ATG12 and HYPK. Densitometric quantification of htt97Qexon1 bands relative to α-tubulin of blots [* P<0.05]. [α-tubulin is loading control in immunoblots. Scale bars in confocal microscopy images represent 5μm. All the presented microscopy and immunoblot data are representative of at least three independent experiments.]

To further investigate if HYPK has specific role in clearance of neddylated proteins through autophagy, we checked the formation and clearance of puromycin-induced neddylated granules in varying knockdown conditions of HYPK, autophagy machinery protein (ATG5) and proteasomal protein (PSMD8). Neddylated protein granules accumulated and persisted even after the puromycin wash in HYPK-KD and ATG5-KD cells **(Figure 7E, 7F).** After puromycin wash, the load of neddylated granules in PSDM8-KD cells decreased to a minimum level that was comparable to control cells **(Figure 7E, 7F).** Additionally, HYPK-KD decreased the conversion of LC3B-I to LC3B-II, which was otherwise observed in Nedd8 overexpressing cells **(Figure 7G).** Such results implied that HYPK functions in the degradation of neddylated proteins through selective autophagy.

Having established the global role of HYPK in neddylation-dependent autophagy, we finally sought to elucidate if HYPK could channelize the degradation of neddylated htt97Qexon1 through aggrephagy. In IMR-32 cells, HYPK colocalized with Nedd8-positive htt97Qexon1-GFP aggregates **(Figure 7H, 7I).** LC3B and Lamp1 also colocalized with the protein complexes of HYPK and htt97Qexon1GFP **(Figure 7H, 7I).** It was also observed that HYPK facilitated the fusion of autophagosomal htt97Qexon1 aggregate with lysosomes to form autolysosomes (AL) **(Figure 6C),** suggesting that the complex of HYPK/neddylated htt97Qexon1-GFP aggregates were not only subjected to be enclosed into autophagic vacuoles, but they were also delivered to lysosomes for degradation. HYPK’s role in autophagic degradation of neddylated htt97Qexon1 was confirmed by following the degradation of the later protein in different conditions that blocked the proteasomal or autophagy pathway by application of specific inhibitors or knockdown of essential proteins of those pathways. HYPK-facilitated degradation of htt97Qexon1 continued in presence of MG132 **(Figure 7J).** However, bafilomycin-A1 treatment of cells had drastically prevented the capacity of HYPK to assist the degradation of htt97Qexon1 **(Figure 7J).** We further validated HYPK’s role as autophagy modulator of htt97Qexon1 in ATG5-KD/HYPK overexpressing cells. Clearance of htt97Qexon1 was impeded in those cells than control cells **(Figure 7K).** Based on these data, we conclude that autophagic degradation of neddylated proteins plays crucial roles in active removal of toxic protein aggregates through the scaffolding functions of HYPK to Nedd8 and LC3 during proteotoxic stress.

## DISCUSSION

The various processes of clearance of intra- and extracellular protein aggregates are tailored to maintain the cellular proteostasis in misfolded protein disorders and neurodegenerative proteinopathies. The complexities in the mechanisms of autophagy are regulated by intriguing activity of different regulatory proteins. In this study, we show that Nedd8 and HYPK are stress-responsive regulators of degradation of protein aggregates by autophagy. The findings suggest that neddylation of cytosolic protein aggregates, and their inclusion in autophagosomes by the receptor function of HYPK **(Figure 8)** are parts of defensive response to control aggrephagy during proteotoxic stress.

**Figure 8.**
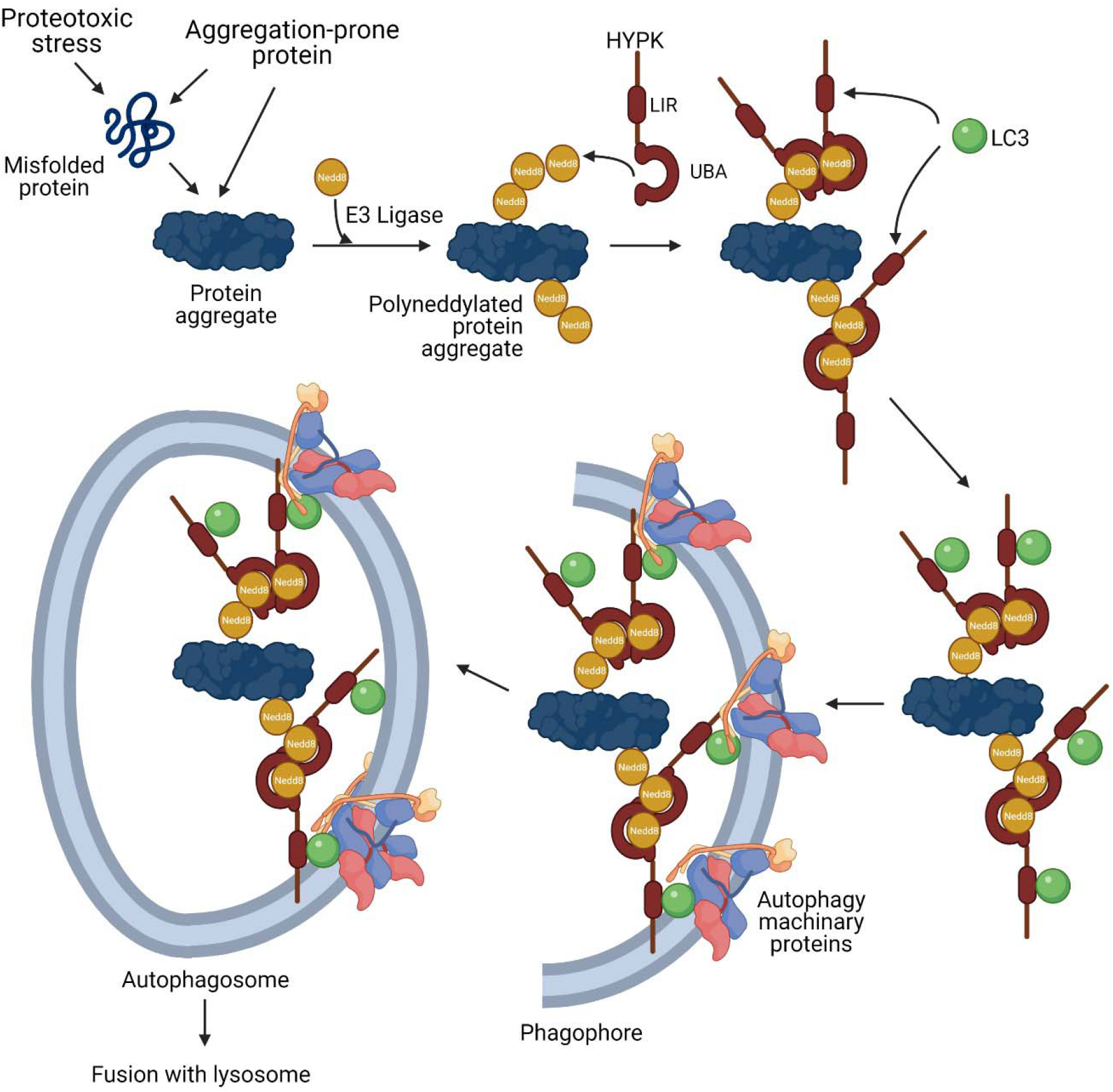
Model of neddylation-dependent aggrephagy and the role of HYPK in this process. In conditions of intrinsic or extrinsic proteotoxic stress, intracellular protein aggregates are progressively neddylated. HYPK binds to the Nedd8 of the polyneddylated protein aggregates by using its C-terminal UBA domain, followed by recruitment of LC3 to the site by HYPK’s LIR. This promotes autophagosomal enclosing of the neddylated protein aggregates for their subsequent degradation upon delivery to lysosome.

In the physiological condition, neddylation has roles in stabilizing the cullin protein of cullin-ring ligase (CRL) complexes (**49**), mitochondrial (**50**) protein etc., other than regulating the activity of ubiquitin ligases (**51**) and maintaining proper localization of ribosomal proteins (**52**). In the stressed conditions, polyneddylation of aggregation-prone proteins leads to their degradation by proteasomal system. Here, we have shown that polyneddylation also prune the aggregated proteins for autophagic degradation in such a way that complements the protein ubiquitination function. We conceive that coexistence of polyubiquitination and polyneddylation pathways in aggrephagy is a part of fail-safe mechanism with far reaching consequences. The differences of ubiquitination and neddylation codes, as defined by the different kinds of lysine linkages in the chains, manifest alternative pathways of protein degradation. While K63-linked polyubiquitin chain delivers proteins to autophagosome, this position-specific lysine is absent in Nedd8. Thus, a unique, yet unknown, kind of neddylation linkage could generate additional topological selectivity towards substrate proteins for autophagy. Additionally, ubiquitination is a global process to most of the cellular proteins, and a large fraction of polyubiquitinated proteins are intended for proteasomal degradation. Contrary to that, neddylation could only be occurring to a limited number of proteins in the direction of aggrephagy. Thus, redundancy of neddylation to such proteins in stressed conditions could compromise their ubiquitination, thereby favouring the autophagic degradation.

Activation-dependent formation of homogeneous or mixed neddylation chains results in differential fate of the substrate proteins. Formation of polyneddylation chain consisting of only Nedd8 protein depends upon the initial charging of the NAE1 (of the NAE1/UBA3 heterodimeric E1 ligase complex) by activated Nedd8 (**53**). The observation that homogeneous polyneddylation of aggregation-prone huntingtin exon1 directs it to autophagic degradation supports the notion that unmixed chains of ubiquitin or Need8 are driving factors in protein degradation pathways, including autophagy. On the other hand, stress-induced formation of mixed ubiquitin-Nedd8 chain results due to UBA1-dependent incorporation of Nedd8 in polyubiquitin chain. Such ubiquitination system-induced Nedd8 addition to ubiquitin chain has an effect in further aggregation of proteins, although that phenomenon still participates in transient protection of cells from cytotoxicity (**20, 54**).

HYPK regulates aggrephagy through its interaction with LC3 and Nedd8, defining the pivotal receptor function of this protein in a noncanonical autophagy pathway. The charged and hydrophobic residues in the UBA domain of the C-terminus are necessary for HYPK binding to Nedd8. The UBA domain is conserved in the longest isoform of HYPK protein of all species, signifying that Nedd8 binding is an essential function of HYPK. Although it is unknown what type lysine-linkage in the polyneddylation chain specifies the autophagosomal degradation of proteins, it is reasonable to speculate that the unique sequence of HYPK-UBA domain is the determinant for recognizing such neddylation linkage. While HYPK recognizes the polyneddylation chain on the protein aggregates, we cannot exclude that it also binds to monomeric Nedd8 and neddylation chain of soluble proteins.

The LC3 binding function is attributed to the N-terminal LIR of HYPK. HYPK has an unconventional LIR core sequence that lacks the hydrophobic amino acid at the fourth position. Although the LIR of HYPK can be categorized as a ‘composite LIR’ (**55**), more structural studies can expand the understanding if it can also function as ‘half LIR’ (**56**). Nevertheless, our preliminary studies found that HYPK interacts with the LIR docking site (LDS) of LC3B, as LDS mutants of LC3B do not interact with HYPK (data not shown). The absence of hydrophobic interactions at the fourth position of HYPK-LIR could be compensated by an extended LIR that is inclusive of other hydrophobic interactions involving not only the HP1 region but also the HP0 and HP2 regions of LC3 and GABARAP proteins. Consistent with our previous report that the LIR-harboring N-terminal region of HYPK is a part of flexible and stretchable disordered nanostructure (**29**), rearrangements of the essential interactions between HYPK-LIR and LDS are possible. While such bonding rearrangements can lower the affinity of HYPK for LC3, oligomeric HYPK binds multiple LC3 molecules, thereby increasing the overall binding strength. Moreover, the LIR is present in the all the isoforms of HYPK, including the splice variant (isoform 2 - NP_001186814.1) that lacks the UBA domain, suggesting that the LC3 binding and autophagy regulatory functions of HYPK are conserved.

The ability of HYPK to regulate neddylation-dependent autophagy links this protein to the general mechanisms of proteostasis. A workable model consists the proposition that proteotoxic stress causes the formation of protein aggregates which are successively neddylated. HYPK interacts with the Nedd8 of the neddylated protein aggregates, followed by sequestration of more HYPK with the aggregates due to its cooperative self-association. Binding with Nedd8 also possibly results in HYPK’s transition from closed-to-open structure due to the loss of intramolecular interactions between the N-terminal charged region and C-terminal low complexity region. The open conformer of HYPK recruits LC3 to the aggresomes, followed by assembly of downstream proteins of autophagy complex for initiation of autophagosomal enclosure of protein aggregates.

Aggrephagy is a major mode of alleviating the load of the pathological protein aggregates in cells of neurodegenerative disorders (NDs) (**57**). Our approach of a global screening of UBLs and UBA domain-containing proteins has identified and validated the functions of Nedd8 and HYPK in modulation of aggrephagy of huntingtin protein. Nedd8 and its conjugation machinery proteins have been identified as modulators of NDs-associated proteins (**58, 59**). NUB1-assited recognistion of mutant huntingtin protein serves as the activation signal for proteasomal degradation of huntingtin protein. Besides the proteasomal pathway, our study showing that genetic and chemical perturbation of neddylation pathway inhibits the aggrephagy of insoluble mutant huntingtin exon1 aggregates strongly suggests that neddylation also causes recruitment of autophagosomal proteins to aggregates by generating Nedd8-HYPK-LC3 complex. The constitutive neddylation of mutant huntingtin could be a segment of bipartite surveillance system for simultaneous and dispensable degradation of the protein by different pathways. The neddylation signal initially primes the soluble mutant huntingtin to proteasome, followed by autophagosomal removal of insoluble aggregates of the protein. It is reported that NUB1 expression, but not the NAE1 expression, is downregulated in knock-in mouse model of Huntington’s disease (HD) (GEO: GDS2912,) (**60**). Thus, it is interesting to test whether autophagy is the primary mode of degradation of neddylated mutant huntingtin, although possibility of other supplementary mechanisms is also obvious. This information can also be extrapolated to speculate that neddylation is prerequisite for autophagic degradation of other labile proteins that are prone to misfolding and aggregation.

Similarly, the spatio-functional aspects of HYPK in proteostasis extend beyond its chaperone (**61**) and aggregate-sequestering (**27**) functions to nonredundant role in huntingtin aggrephagy. Although HYPK reportedly exerts its chaperone activity by interacting with the N-terminal region of huntingtin (**62**), we reveal that aggrephagy is the mechanism of HYPK-mediated removal of huntingtin aggregates. While the coupling of mutant huntingtin, HYPK, Nedd8 and LC3 can be dependent upon the context of layers of huntingtin aggregates, it is also interesting to note that the expression of HYPK is regulated by an array of factors like thermal stress (**63**), cellular aging (**33**) etc. Our previous studies showed that not only huntingtin, but other disease-causing aggregation-prone proteins also elicit a HYPK coaggregation response (**27**). Furthermore, the recruitment of HYPK to neddylated proteins can also be remotely governed by the acetylation status of the target protein. The interaction of HYPK with the proteins of N-acetyl transferase (NAT) complex (**64**) and acetylation-dependent neddylation of proteins (**65**) are indicative of a complex crosstalk of post-translational modifications and HYPK recruitment in multiprotein complexes during aggrephagy of misfolded proteins. Overall, the neddylation-dependent aggrephagy takes place by synergistic effects of sequestration of HYPK with protein aggregates and tethering function of HYPK.

Information of expression profile derived from gene expression omnibus (GEO) show that HYPK expression is not significantly changed in different NDs and brain diseases **(Supporting table 1**). Many of the HYPK interacting proteins also contain one or more aggregation-prone regions (data not shown), suggesting that proteostatic surveillance is general function of HYPK. Although HYPK is not a druggable target due to its noncatalytic function, modulation of its expression could be a potential mean to cope with the challenges of ND-associated protein aggregates.

In summary, this study identified the functionality of Nedd8 and HYPK as the modulators of autophagy in context of clearance of proteotoxic stress-induced and innate protein aggregates, thereby describing a novel and noncanonical mechanism of aggrephagy that can be utilized in HD therapeutics.

## ACKNOWLEDGEMENTS

The authors extend their thankfulness to the Biocluster of National Centre for Biological Sciences (India) for providing the TEM facility. The authors also thank the members of CFG group for giving critical inputs in different experiments and manuscript writing. The personnel of sophisticated equipment facility of the research support service group of CDFD are acknowledged for the technical assistance in operating different instruments.

## AUTHOR CONTRIBUTIONS

DKG: Conceptualization of the rationale of study, performed experiments, data analysis, manuscript writing. AR: Conceptualization of the rationale of study, data analysis, arrangement of funds, mentoring.

## MATERIALS and METHODS

### Clones and plasmids

To clone the open reading frames (ORFs) of different genes, total RNA was isolated (ThermoFischer Scientific [TFS], 12183018A) from IMR-32 cells, and cDNAs of corresponding mRNAs were made by reverse transcription using the oligo-dT primer (Eurofins Genomics [EG]). ORFs of genes were amplified from cDNAs by polymerase chain reaction (PCR) using specific primer sets (EG). The list of clones and plasmids used in this study is given in **Table 1.** The general process of cloning was similar to what was described in our previous studies (**27, 66**). Restriction digested PCR products and plasmids were ligated, followed by the transformation of the ligated products in ultra-competent DH5α strain of *Escherichia coli.* Positive clones were selected by colony PCR. All clones were sequenced at the sophisticated equipment facility of the research support service group of CDFD.

**Table 1.**
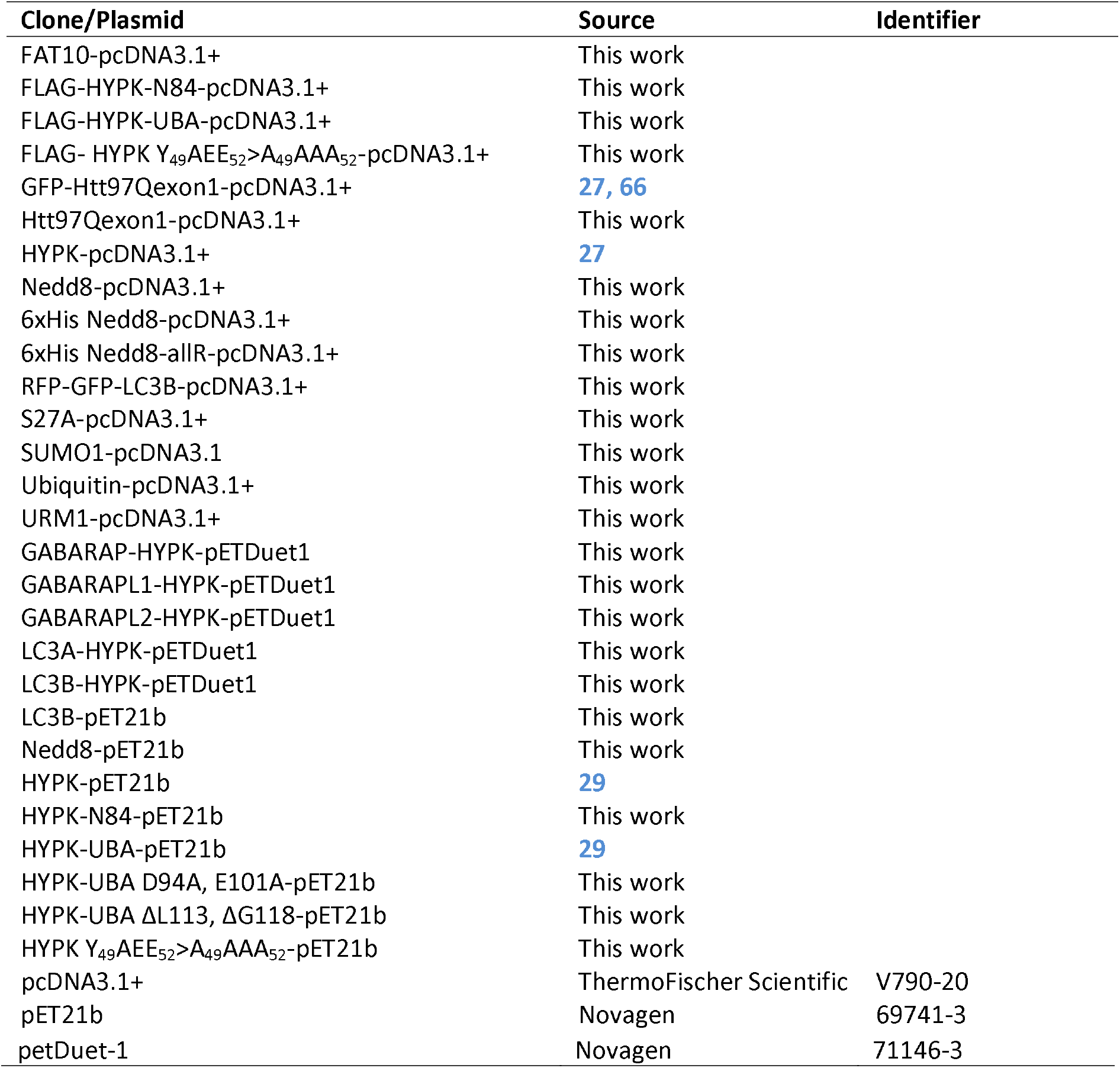
Clones and plasmids

### Site-directed mutagenesis

Deletion mutants of HYPK were made by using specific primer sets (EG) in PCR-based method. FLAG peptide sequence was introduced to different clones by incorporating the sequence in-frame at the N-terminus of ORFs by using the forward primer (EG). Point mutations in HYPK (HYPK Y_49_AEE_52_>A_49_AAA_52_), UBA domain of HYPK (UBA D94A, E101A; UBA ΔL113, ΔG118) and Nedd8 (allR - all lysine-to-arginine) were created by using multiple primer sets (EG) in overlapping PCR method. Mutations were confirmed by sequencing.

### Recombinant protein production and purification

Recombinant HYPK, HYPK-UBA, HYPK-UBA D94A E101A, HYPK-UBA ΔL113 ΔG118, HYPK-N84, HYPK Y_49_AEE_52_>A_49_AAA_52_, Nedd8, LC3A, LC3B, GABARAP, GABARAPL1, GABARAPL2 proteins were produced by using a T7 expression system that was described by us in earlier studies (**29, 67**). The bacterial expression clones (in pET21 and pETDuet-1 vectors) were separately transformed in the BL21DE3 strain of *Escherichia coli*, followed by induction of protein synthesis by application of isopropyl-β-D-1-thiogalactopyranoside (IPTG, 1mM final concentration) (SA, I6758) in the culture medium (Luria Bertani broth/ampicillin). After 12-18 hours of protein production at 1°C/180rpm, bacterial cells were lysed by sonication in the lysis buffer [50mM Tris-Cl (pH: 8.0), 300mM NaCl, 10mM imidazole, 1mM PMSF], Cell lysate was cleared by centrifugation (14000g/40 minutes/4°C), followed by purification of proteins by nickel ion-based affinity exchange column chromatography. The cleared lysate was run through Ni^+2^-NTA agarose beads (Qiagen [QA], 30250) to allow binding of the histidine-tagged (recombinant) proteins to beads. The beads were subsequently washed with wash buffer [50mM Tris-Cl (pH: 8.0), 300mM NaCl, 40mM imidazole], followed by elution of the proteins in elution buffer [50mM Tris-Cl (pH: 8.0), 300mM NaCl, 300mM imidazole]. Proteins were dialyzed in different dialysis buffers depending upon the requirements of downstream experiments. In all the experiments, >98% pure proteins were used.

### Protein-protein binding assay by surface plasmon resonance

The binding properties of HYPK, HYPK-UBA, HYPK-UBA D94A, E101A, HYPK-UBA ΔL113, ΔG118 with Nedd8; and HYPK, HYPK-N84, HYPK-UBA, HYPK Y_49_AEE_52_>A_49_AAA_52_ with LC3B were analyzed by surface plasmon resonance (SPR) technique. Using the NHS/EDC reagent, 5nmole of Nedd8 or LC3B [in acetate buffer (pH: 4.0)] was immobilized on a CM5 sensor chip (GE Healthcare Life Sciences, 29149604) by amine coupling. The mobile phase analytes (HYPK, HYPK-UBA, HYPK-UBA D94A, E101A and HYPK-UBA ΔL113, ΔG118, HYPK-N84, HYPK Y_49_AEE_52_>A_49_AAA_52_) were kept in HBS-EP buffer [10mM HEPES (pH: 7.5), 150mM NaCl, 3mM EDTA, 0.005% (v/v) Surfactant P20 (pH: 7.4)]. Analyte to ligand binding experiments were done in the Biacore 3000 instrument (GE Healthcare Life Sciences) at 25°C. Analytes were injected at a flowrate of 30μl/min, and the dissociation events were allowed for 10 minutes. Concentrations of analytes were in the range of 40nM-25μM. Subtraction of nonspecific binding from the actual binding response was done by measuring the mock binding on the immobilized surface of another channel. Dissociation constants (K_d_) were generated by assuming 1:1 Langmuir model of interaction.

### Cell culture and reagent treatment

The MCF7 and IMR-32 cell-lines were procured from National Centre for Cell Sciences (NCCS, India). MCF7 cells were cultured in advanced DMEM medium (TFS, 12491023) that was supplemented with 10% fetal bovine serum (TFS, 26140095), 2mM L-glutamine (TFS, 25030164) and 1x antibiotic-antimycotic solution (TFS, 11548876). IMR-32 cells were grown in neuronal culture medium (TFS, 88283) supplemented with other components as mentioned for MCF7 cells. Cells were maintained in a humified incubator at 37°C with 5% CO_2_. Although cell lines were not authenticated, they were routinely checked for mycoplasma contamination by using Venor™ GeM mycoplasma detection kit [Sigma-Aldrich (SA), MP0025]. The stable expression of RFP-GFP-LC3B (in pcDNA3.1÷) in MCF7 cell line, and GFP-htt97Qexon1 (in pcDNA3.1+) or htt97Qexon1 (in pcDNA3.1+) in IMR-32 cell-line were selected by keeping 2μM neomycin (SA, PHR1491) in the cell culture medium.

Puromycin (SA, P8833), MLN4924 [SA (Calbiochem), 505477), MG132 [SA (Calbiochem), 474790], Bafilomycin-A1 (SA, 196000), and MLN7243 (Aobious, AOB87172) were treated to cells with specific concentration for defined time (as mentioned in different sections of results).

Clones and plasmids were transfected to subconfluent cells by using Lipofectamine 2000 (TFS, 11668019) and Opti-MEM (TFS, 31985070) medium following manufacturer’s instructions. siRNAs were transfected to cells by using Lipofectamine 3000 (TFS, L3000015). List of siRNAs are given in **Table 2.** Transfected cells were harvested at different timepoints after transfection depending upon the experimental requirements. In chase experiments, cells were intermittently harvested at an interval of 6 or 12 hours.

**Table 2.** esiRNAs and siRNAs

| Target of esiRNA/siRNA | Source | Identifier |
| --- | --- | --- |
| ATG5 | Sigma-Aldrich | EHU085781 |
| CBL | Sigma-Aldrich | EHU032851 |
| CBL-B | Sigma-Aldrich | EHU074901 |
| FAT10 | Sigma-Aldrich | EHU230651 |
| HYPK | Sigma-Aldrich | EHU096201 |
| HUWE1 | Sigma-Aldrich | EHU067731 |
| MARK1 | Sigma-Aldrich | EHU075901 |
| MARK2 | Sigma-Aldrich | EHU134171 |
| MARK3 | Sigma-Aldrich | EHU093021 |
| MARK4 | Sigma-Aldrich | EHU903821 |
| Nedd8 | Sigma-Aldrich | EHU112521 |
| NUB1 | Sigma-Aldrich | EHU020291 |
| PSMD8 | Sigma-Aldrich | EHU147871 |
| Rad23A | Sigma-Aldrich | EHU027621 |
| Rad23B | Sigma-Aldrich | EHU145881 |
| RSC1A1 | Sigma-Aldrich | EHU230881 |
| S27A | Sigma-Aldrich | EHU112411 |
| SIK1 | Sigma-Aldrich | EHU138711 |
| SQSTM1 | Sigma-Aldrich | EHU027651 |
| SUMO1 | Sigma-Aldrich | EHU106621 |
| TDRD3 | Sigma-Aldrich | EHU037161 |
| TNRC6C | Sigma-Aldrich | EHU080181 |
| UBAC1 | Sigma-Aldrich | EHU143751 |
| UBAC2 | Sigma-Aldrich | EHU141521 |
| UBAP2 | Sigma-Aldrich | EHU079831 |
| UBE2K | Sigma-Aldrich | EHU091111 |
| Ubiquitin | Santa Cruz Biotechnology | sc-29513 |
| UBL7 | Sigma-Aldrich | EHU060951 |
| UBP5 | Sigma-Aldrich | EHU025811 |
| UBP13 | Sigma-Aldrich | EHU003961 |
| UBQL1 | Sigma-Aldrich | EHU084751 |
| UBQL2 | Sigma-Aldrich | EHU029491 |
| UBQL3 | Sigma-Aldrich | EHU044731 |
| UBQL4 | Sigma-Aldrich | EHU109861 |
| UBS3B | Sigma-Aldrich | EHU008351 |
| UBXN1 | Sigma-Aldrich | EHU157321 |
| URM1 | Sigma-Aldrich | EHU009021 |
| VPS13D | Sigma-Aldrich | EHU123621 |
| Universal negative control | Sigma-Aldrich | SIC001 |

### Fractionation of soluble proteins and insoluble protein aggregates from cell lysate

To separate the insoluble polyneddylated protein aggregates and/or htt97Qexon1 aggregates from the soluble proteins, cells were scrapped from cell culture dishes in phosphate buffer saline (PBS, pH: 7.4), followed by lysis of cells in pre-chilled cell lysis buffer [LB - 50mM Tris-Cl (pH: 8.0), 50mM NaCl, 5mM MgCl_2_, 0.2% Triton X-100, 0.5% Nonidet P-40, 1x protease inhibitor cocktail [SA, 11836170001)], phosphatase inhibitor (SA, PHOSS-RO)]. After keeping the cell suspension at 4°C for 15 minutes in LB, cell lysate was cleared by centrifugation at 14000g for 30 minutes (4°C). The supernatant contained the fraction of soluble proteins (SF). The precipitate, which contained insoluble fraction (IF) of protein aggregates and other cell debris, was resolubilized in LB that was supplemented with 1% SDS and 4M guanidinium chloride, followed by centrifugation of the suspension at 14000g for 30 minutes (4°C). The supernatant, which contained high-molecular weight polyneddylated proteins and/or htt97Qexon1 proteins, was finally dialyzed in dialysis buffer [DB – 50mM Tris-Cl (pH: 8.0), 50mM NaCl].

### Immunoprecipitation and Immunoblotting

Immunoprecipitation (IP) of desired proteins were done from SF (2mg of protein) or SF (1mg of protein) + IF (1mg of protein) by using crosslink magnetic IP/co-IP method [TFS (Pierce Crosslink Magnetic IP/co-IP kit), 88805) following the manufacturer’s protocol. Some IP experiments were done in denaturing conditions (with 1% SDS) as mentioned in the result section. List of antibodies used in IPs is given in **Table 3.**

**Table 3.** Antibodies

| Primary antibodies |  |  |  |
| --- | --- | --- | --- |
| Target protein | Species | Supplier | Catalogue number |
| Alpha-tubulin | Mouse | Sigma-Aldrich | T9026 |
| ATG5 | Rabbit | Sigma-Aldrich | A0856 |
| BECN1 | Rabbit | Cell Signalling Technology | 3738 |
| Beta-tubulin | Mouse | Sigma-Aldrich | T8328 |
| FAT10 | Mouse | Abcam | ab168680 |
| FLAG | Mouse | Sigma-Aldrich | F3165 |
| His-tag | Mouse | Sigma-Aldrich | SAB2702218 |
| HYPK | Rabbit | Sigma-Aldrich | SAB1101781 |
| Lamp1 | Mouse | Abcam | ab25630 |
| LC3B | Rabbit | Abcam | ab48394 |
| LC3B | Mouse | Cell Signalling Technology | 83506 |
| Nedd8 | Rabbit | Abcam | ab194582 |
| Nedd8 | Mouse | ThermoFischer Scientific | MA5-17133 |
| p62/SQSTM1 | Mouse | Abcam | ab56416 |
| polyQ (65Q) | Mouse | Sigma-Aldrich | P1874 |
| pS15-BECN1 | Rabbit | Cell Signalling Technology | 84966S |
| S27A | Rabbit | Abcam | ab227011 |
| SUMO1 | Rabbit | Abcam | ab227424 |
| Ubiquitin | Rabbit | Abcam | ab223613 |
| URM1 | Rabbit | Abcam | ab220490 |
| Secondary antibodies<br>(Immunoblotting) |  |  |  |
| anti-rabbit IgG (whole molecule) peroxidase antibody | Goat | Sigma-Aldrich | A0545 |
| anti-mouse IgG (whole molecule) peroxidase antibody | Rabbit | Sigma-Aldrich | A9044 |
| Secondary antibodies<br>(Immunocytochemistry) |  |  |  |
| anti-Mouse IgG (H+L)-Alexa Fluor Plus 488 | Goat | ThermoFischer Scientific | A32723 |
| anti-Mouse IgG (H+L)-Alexa Fluor Plus 555 | Goat | ThermoFischer Scientific | A32727 |
| anti-Mouse IgG (H+L)-<br>Alexa Fluor Plus 647 | Goat | ThermoFischer Scientific | A32728 |
| anti-Rabbit IgG (H+L)-<br>Alexa Fluor Plus 488 | Goat | ThermoFischer Scientific | A32731 |
| Anti-Rabbit IgG (H+L)-<br>Alexa Fluor Plus 555 | Goat | ThermoFischer Scientific | A32732 |
| anti-Rabbit IgG (H+L)-<br>Alexa Fluor Plus 647 | Goat | ThermoFischer Scientific | A32733 |

In immunoblotting (IB), 60-80μg of total protein was added with 4x Laemmli buffer (Bio-Rad, 161-0747) and heated at 95°C for 10 minutes. Proteins were separated on 12% SDS-polyacrylamide gel, followed by transfer of proteins from gel to polyvinylidene fluoride (PVDF) membrane (Amersham Hybond P 0.45, GE Healthcare Life Sciences) and sequential use of primary and secondary antibodies that are listed in **Table 3.** Intermittent washing was done by TBST buffer (pH: 7.4). Chemiluminescent signal was generated by SuperSignal west femto maximum sensitivity substrate (TFS, 34094) and detected by ChemiDoc XRS+ imaging system (Bio-Rad). Densitometric quantification of bands was done in ImageJ2 software (**68**).

### Immunocytochemistry and fluorescence microscopy

MCF7 or IMR-32 cells were grown on glass coverslips. The adherent cells were washed with PBS, followed by fixation with methanol or 4% paraformaldehyde (in PBS). Cell permeabilization was done with 0.2% TritonX-100 (in PBS, pH: 7.4), followed by blocking with 1% BSA (in PBS). Primary and secondary antibodies (listed in **Table 3)** were sequentially given to cells with intermittent washing with PBS. Cells were mounted in prolong antifade gold with or without DAPI (TFS, P36931; P10144). Images were acquired in Zeiss LSM700 confocal laser scanning microscope (Carl Zeiss) with the 63x Plan-Apochromat/ 1.4 NA oil/ DIC M27 objective. Images were processed in Zen-Lite 2010 software (Carl Zeiss). Colocalization analysis of different proteins was done in Zen-Lite 2010 (blue edition) software (Carl Zeiss) by using the colocalization algorithm. The Pearson’s correlation coefficient was used as a measure of colocalization of proteins.

The LC3B-positive autophagosome puncta, Nedd8 granules and htt97Qexon1-GFP aggregates in cells were counted in Eclips Ti E (Nikon) microscope by using NIS-elements imaging software.

### Autophagy flux assay

The GFP+/RFP+ and GFP-/RFP+ LC3B puncta represented the autophagosomes and autolysosomes respectively, and their numbers were used to measure the autophagy flux as mentioned in previous study (**69**). To monitor the autophagy flux in varying knockdown condition of different proteins, cells were transfected with 10nM of esiRNA against target mRNA. The culture medium was changed after 48 hours of transfection.

### Transmission electron microscopy

IMR-32 cells were grown in culture dishes. HYPK or htt97Qexon1+HYPK were transfected to cells by the procedure that is mentioned in cell culture section. Cells were detached from dishes by trypsin-EDTA solution (SA, T3924). Detached cells were fixed in 1.5% (v/v) glutaraldehyde/4% (w/v) formaldehyde (in 0.1M cacodylate buffer, pH: 7.4) solution for five hours, followed by washing of cells with PBS (pH: 7.4) for few times. Dehydration of the cells was gradually done by sequentially keeping them in 50%, 70% and 90% ethanol (15 minutes/solution). Cells were sectioned (1μm thick) in ultra-microtome before placing them on 200 mesh copper grid (SA, G4776). Cell-section containing grids were incubated in 0.05M glycine (in PBS) for 20 minutes. Grids were then washed with PBS, followed by staining with aqueous uranyl acetate for 2 minutes. After the grids were washed with water, they were transiently (for 15 seconds) exposed to lead citrate. Images were collected in Tecnai G2 Spirit Bio-TWIN transmission electron microscope. Electron beam strength was 15kV and images were recorded in Gatan Orins CCD camera.

### Computational studies

#### Protein-protein docking

Structures of Nedd8 protein and the UBA domains of different human proteins were curated (listed in **Table 4)** from the Protein Data Bank (**70**). Structures of the proteins were optimized to add missing hydrogen atoms and assign bond orders in the protein preparation wizard of Maestro 9.2 (Schrödinger) [forcefield: OPLS_2005, convergence of heavy atoms to root mean square deviation (RMSD): 0.3 Å] as described in our previous studies (**71, 72**). Structures were subsequently minimized in molecular modeling toolkit (**73**) by assisted model building with energy refinement (AMBER) (**74**) simulation to 10^3^ steepest descent iteration. Unrestrained protein-protein docking of the finally prepared structures was done by using the PIPER program (**75**) in the BioLuminate suite (Maestro 9.2, Schrödinger). The stability of complexes was measured by analyzing the ΔG(diss) in the PDBePISA (Proteins, Interfaces, Structures and Assemblies) web server (**76**). Structures were visualized and interpreted in PyMOL.

**Table 4.** PDB identifiers of proteins’ structures

| UBA domain of protein | PDB identifier | Reference |
| --- | --- | --- |
| CBL | 2O09 | 77 |
| CBL-B | 2JNH | 78 |
| HUWE1 | 5LP8 | 79 |
| HYPK | 6C95 | 46 |
| MARK1 | 2HAK | 80 |
| MARK2 | 1Y8G | 81 |
| MARK3 | 2QNJ | 82 |
| NBR1 | 2MGW | 83 |
| Nedd8 | 1NDD | 84 |
| NUB1 | 1WJU | Unpublished |
| Rad23A | 1DV0 | 85 |
| Rad23B | 1P1A | 86 |
| SQSTM1 | 2K0B | 87 |
| TDRD3 | 1WJI | Unpublished |
| TNRC6C | 2DKL | Unpublished |
| UBAC1 | 2DAI | Unpublished |
| UBAP1 | 1WGN | Unpublished |
| UBE2K | 5DFL | 88 |
| Ubiquilin1 | 2JY6 | 89 |
| Ubiquilin3 | 2DAH | Unpublished |
| Ubiquilin4 | 2KNZ | 90 |
| UBL7 | 2CWB | 91 |
| USP5 | 2DAG | Unpublished |

#### Sequence alignment

The sequences of ubiquitin, UBLs and HYPK of different organisms were obtained from protein database of National Center for Biotechnology Information (NCBI). Sequences of UBA domains of different human proteins were curated from the SMART database (**92**). The alignments of sequences were done in Clustal Omega (**93**).

#### Phylogenetic analysis

The sequence alignment file of the UBA domains of different human proteins was saved in the phylip (.phy) format which was further used in the Phylip-3.695 software package. The Seqboot, Protdist, Neighbor and Consense programs were sequentially run to analyze the phylogeny of the UBA domains. The outfile of each program was taken as the input file of the next program. Details of the parameters of each program are available upon request. The cladogram was generated in the Treeview software.

### Statistical analysis

The statistical significance of difference of means between groups were analyzed by two-tailed, homoscedastic student’s t-test. P<0.05 represented statistically significant difference.

### Graphics

The graphics were made in Adobe Illustrator.

## DATA AVAILABILITY

All the experimental protocols, data and resources of this study are available from the corresponding authors on request.

## SUPPORTING FIGURES AND LEGENDS

**Supporting figure 1.**
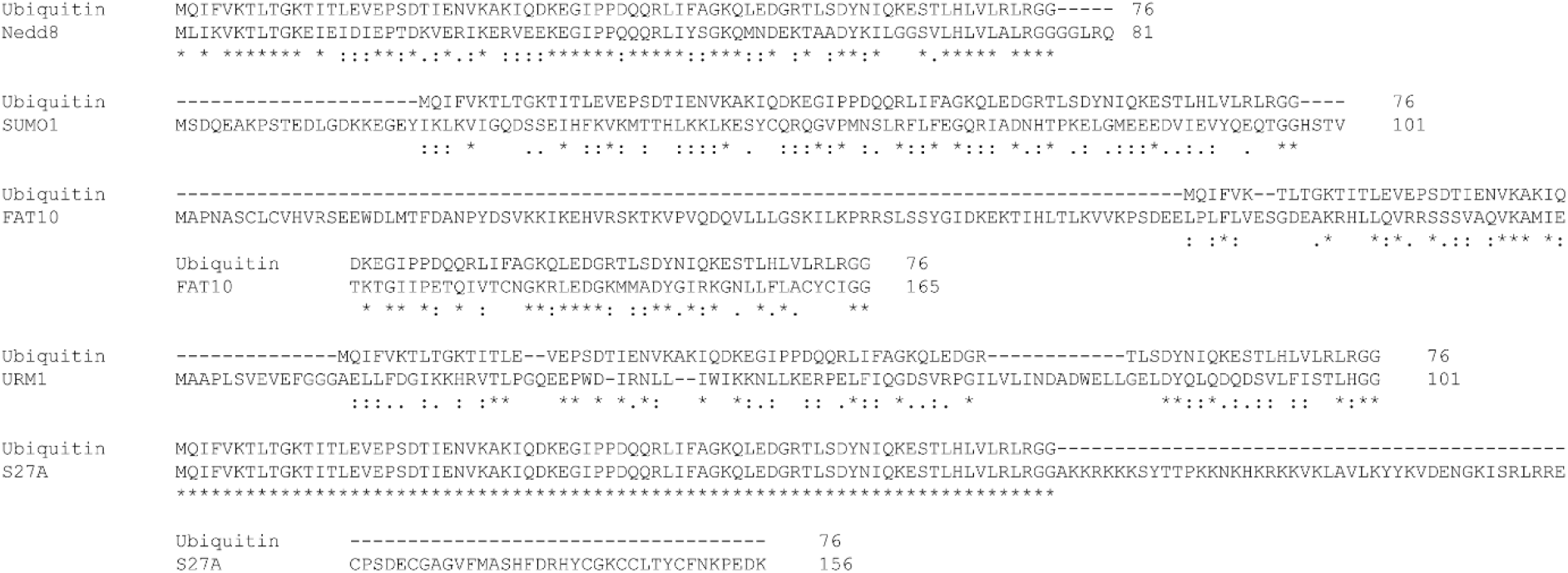
Sequence alignment of ubiquitin with different ubiquitin-like proteins.

**Supporting figure 2.**
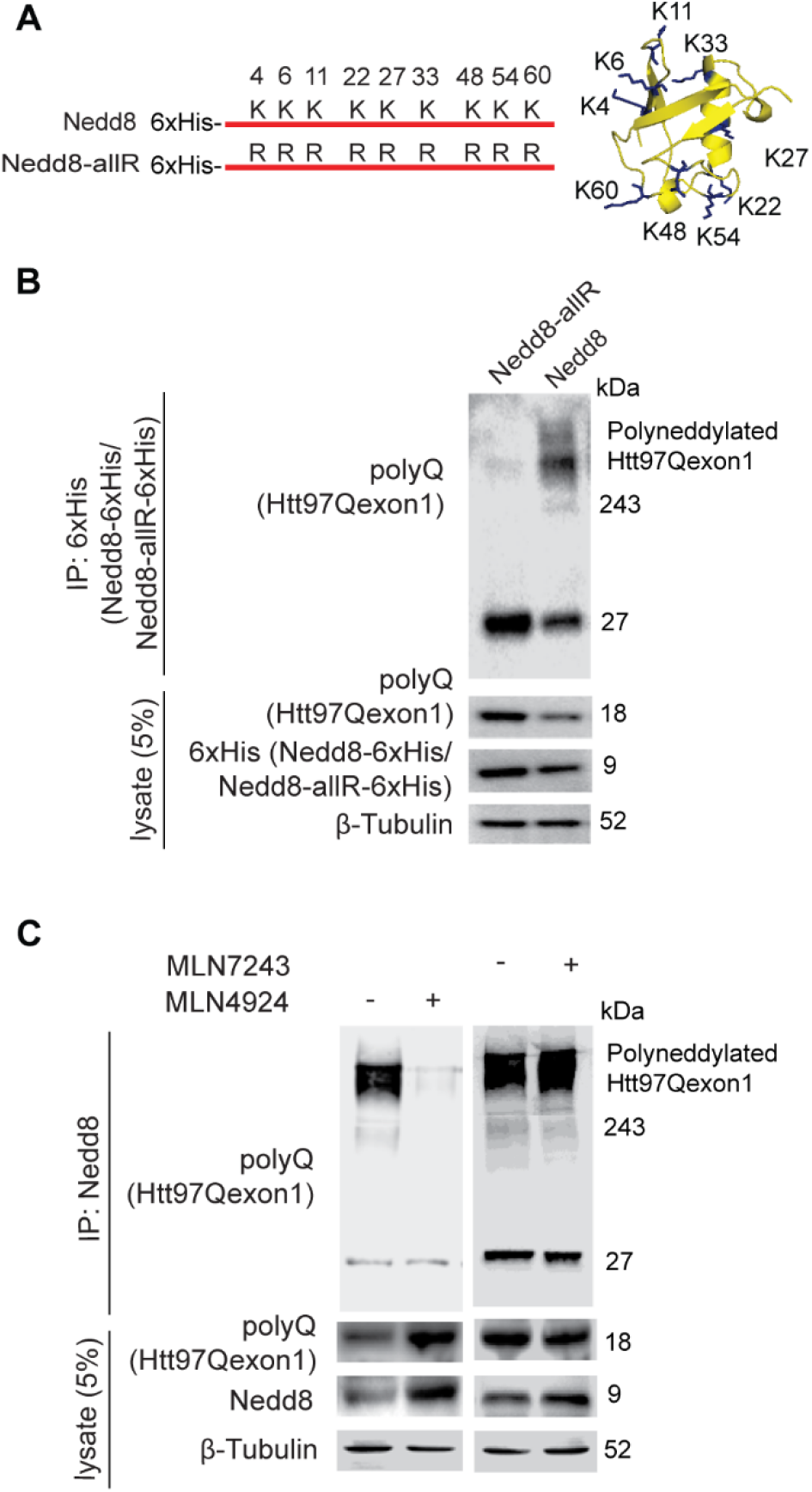
Htt97Qexon1 undergoes homogeneous polyneddylation. (A) Left: schematic representation of wild type and mutant Nedd8 constructs. In the Nedd8-allR mutant, all the lysine residues were arginine. Positions of lysine residues in the wild-type human Nedd8 are indicated. Right: crystal structure of Nedd8 [84] with lysine residues indicated with stick structure. (B) Representative Immunoblots of htt97Qexon1, 6xHis-tagged Nedd8 and 6xHis-tagged Nedd8-allR of immunocomplexes of 6xHis-Nedd8 and 6xHis-Nedd8-allR from lysate of stable htt97Qexon1 expressing IMR-32 cells. (C) Stable htt97Qexon1 expressing IMR-32 cells were treated with 1μM MLN4924 or 1μM MLN7243 for 24 hours. Representative Immunoblots of htt97Qexon1 and Nedd8 of immunocomplexes of Nedd8 from lysate of the above-mentioned cells. [β-tubulin is loading control in immunoblots. Immunoblot data are representative of at least three independent experiments.]

**Supporting figure 3.**
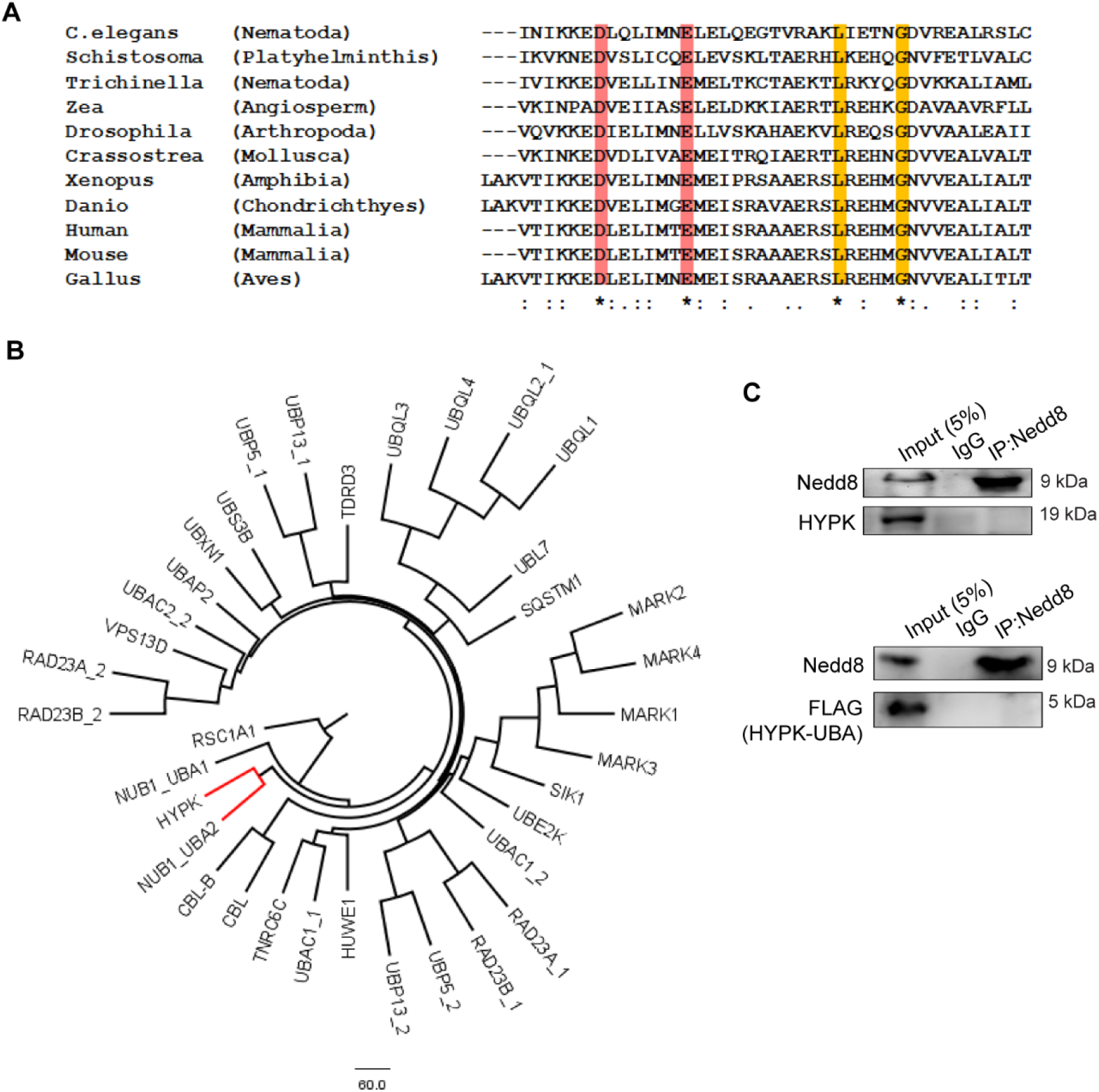
The conserved UBA domain of HYPK noncovalently binds to Nedd8. (A) Multiple sequence alignment of the UBA domains of HYPK proteins of different organisms. Conserved charged and hydrophobic amino acids are highlighted with red and yellow colour respectively. (B) Cladogram representing the clustered UBA domains of different human proteins. Scale indicates phylogenetic distance. (C) Representative immunoblots of Nedd8, HYPK and FLAG-tagged HYPK-UBA [ectopically expressed] of denaturing immunoprecipitation of Nedd8 from lysate of MCF7 cells. [Immunoblot data are representative of at least three independent experiments.]

**Supporting table 1.**
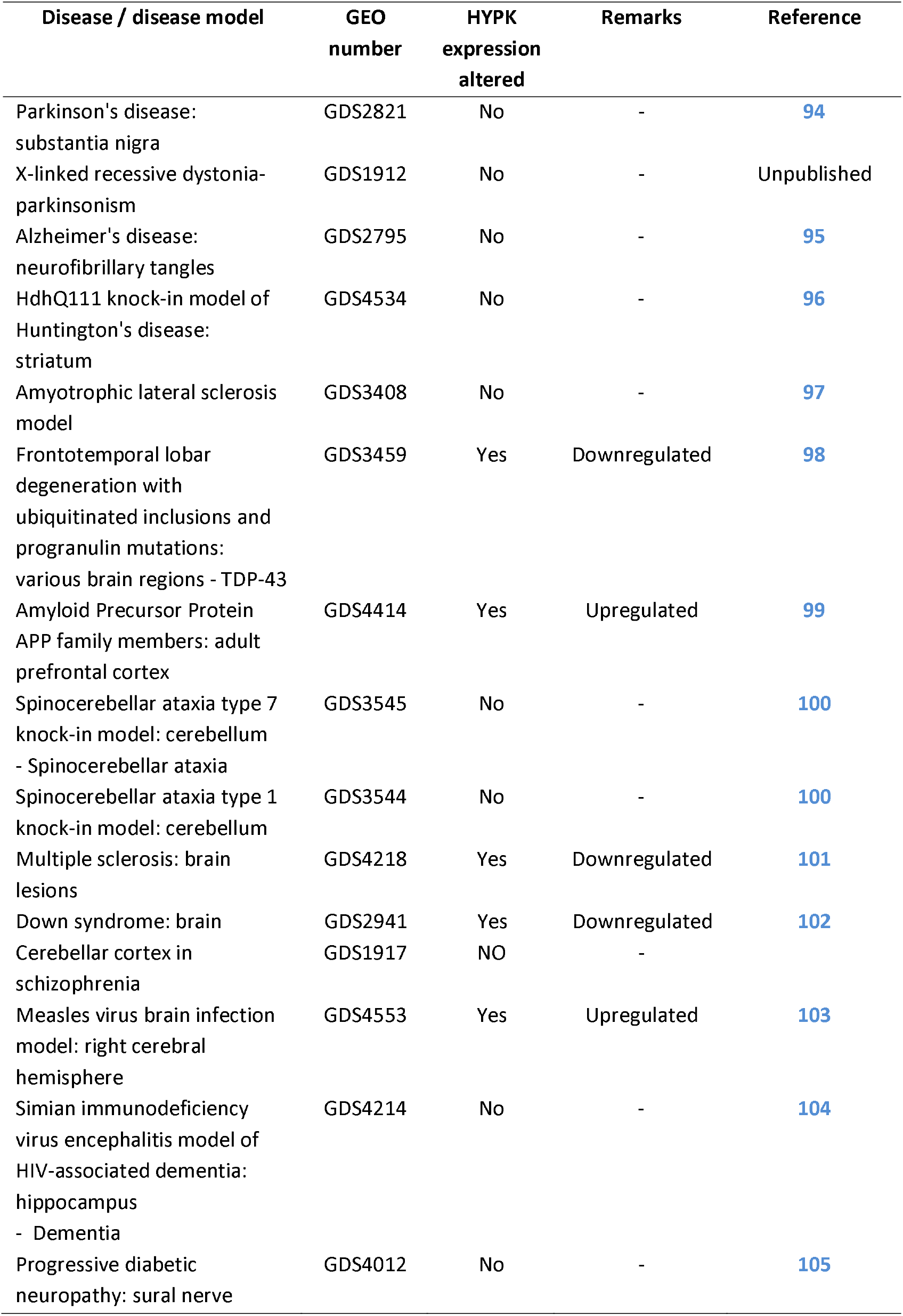
HYPK transcript expression level in different neurodegenerative and brain diseases.

## REFERENCES

1. Labbadia, J. & Morimoto, R. I. The biology of proteostasis in aging and disease. Annu Rev Biochem 84, 435–464, doi:10.1146/annurev-biochem-060614-033955 (2015).

2. Klaips, C. L., Jayaraj, G. G. & Hartl, F. U. Pathways of cellular proteostasis in aging and disease. J Cell Biol 217, 51–63, doi:10.1083/jcb.201709072 (2018).

3. Chen, B., Retzlaff, M., Roos, T. & Frydman, J. Cellular strategies of protein quality control. Cold Spring Harb Perspect Biol 3, a004374, doi:10.1101/cshperspect.a004374 (2011).

4. Kim, Y. E., Hipp, M. S., Bracher, A., Hayer-Hartl, M. & Hartl, F. U. Molecular chaperone functions in protein folding and proteostasis. Annu Rev Biochem 82, 323–355, doi:10.1146/annurev-biochem-060208-092442 (2013).

5. Tanaka, K. The proteasome: overview of structure and functions. Proc Jpn Acad Ser B Phys Biol Sci 85, 12–36, doi:10.2183/pjab.85.12 (2009).

6. Glick, D., Barth, S. & Macleod, K. F. Autophagy: cellular and molecular mechanisms. J Pathol 221, 3–12, doi:10.1002/path.2697 (2010).

7. Lamark, T. & Johansen, T. Aggrephagy: selective disposal of protein aggregates by macroautophagy. Int J Cell Biol 2012, 736905, doi:10.1155/2012/736905 (2012).

8. Mizushima, N. Autophagy: process and function. Genes Dev 21, 2861–2873, doi:10.1101/gad.1599207 (2007).

9. Hurley, J. H. & Young, L. N. Mechanisms of Autophagy Initiation. Annu Rev Biochem 86, 225–244, doi:10.1146/annurev-biochem-061516-044820 (2017).

10. Wesch, N., Kirkin, V. & Rogov, V. V. Atg8-Family Proteins-Structural Features and Molecular Interactions in Autophagy and Beyond. Cells 9,doi:10.3390/cells9092008 (2020).

11. Ruz, C., Alcantud, J. L., Vives Montero, F., Duran, R. & Bandres-Ciga, S. Proteotoxicity and Neurodegenerative Diseases. Int J Mol Sci 21,doi:10.3390/ijms21165646 (2020).

12. Chen, R. H., Chen, Y. H. & Huang, T. Y. Ubiquitin-mediated regulation of autophagy. J Biomed Sci 26, 80, doi:10.1186/s12929-019-0569-y (2019).

13. Zheng, N. & Shabek, N. Ubiquitin Ligases: Structure, Function, and Regulation. Annu Rev Biochem 86, 129–157, doi:10.1146/annurev-biochem-060815-014922 (2017).

14. Tan, J. M. et al. Lysine 63-linked ubiquitination promotes the formation and autophagic clearance of protein inclusions associated with neurodegenerative diseases. Hum Mol Genet 17, 431–439, doi:10.1093/hmg/ddm320 (2008).

15. Lorente, M. et al. Inhibiting SUMO1-mediated SUMOylation induces autophagy-mediated cancer cell death and reduces tumour cell invasion via RAC1. J Cell Sci 132,doi:10.1242/jcs.234120 (2019).

16. Enchev, R. I., Schulman, B. A. & Peter, M. Protein neddylation: beyond cullin-RING ligases. Nat Rev Mol Cell Biol 16, 30–44, doi:10.1038/nrm3919 (2015).

17. Abidi, N. & Xirodimas, D. P. Regulation of cancer-related pathways by protein NEDDylation and strategies for the use of NEDD8 inhibitors in the clinic. Endocr Relat Cancer 22, T55–70, doi:10.1530/ERC-14-0315 (2015).

18. Jayabalan, A. K. et al. NEDDylation promotes stress granule assembly. Nat Commun 7, 12125, doi:10.1038/ncomms12125 (2016).

19. Mori, F. et al. Accumulation of NEDD8 in neuronal and glial inclusions of neurodegenerative disorders. Neuropathol Appl Neurobiol 31, 53–61, doi:10.1111/j.1365-2990.2004.00603.x (2005).

20. Maghames, C. M. et al. NEDDylation promotes nuclear protein aggregation and protects the Ubiquitin Proteasome System upon proteotoxic stress. Nat Commun 9, 4376, doi:10.1038/s41467-018-06365-0 (2018).

21. Lane, D. P. Stress, specificity and the NEDD8 proteome. Cell Cycle 11, 1488–1489, doi:10.4161/cc.20073 (2012).

22. Li, J. et al. NEDD8 Ultimate Buster 1 Long (NUB1L) Protein Suppresses Atypical Neddylation and Promotes the Proteasomal Degradation of Misfolded Proteins. J Biol Chem 290, 23850–23862, doi:10.1074/jbc.M115.664375 (2015).

23. Behrends, C. & Fulda, S. Receptor proteins in selective autophagy. Int J Cell Biol 2012, 673290, doi:10.1155/2012/673290 (2012).

24. Lippai, M. & Low, P. The role of the selective adaptor p62 and ubiquitin-like proteins in autophagy. Biomed Res Int 2014, 832704, doi:10.1155/2014/832704 (2014).

25. Sarraf, S. A. et al. Loss of TAX1BP1-Directed Autophagy Results in Protein Aggregate Accumulation in the Brain. Mol Cell 80, 779–795 e710, doi:10.1016/j.molcel.2020.10.041 (2020).

26. Lu, K., Psakhye, I. & Jentsch, S. Autophagic clearance of polyQ proteins mediated by ubiquitin-Atg8 adaptors of the conserved CUET protein family. Cell 158, 549–563, doi:10.1016/j.cell.2014.05.048 (2014).

27. Ghosh, D. K., Roy, A. & Ranjan, A. Aggregation-prone Regions in HYPK Help It to Form Sequestration Complex for Toxic Protein Aggregates. J Mol Biol 430, 963–986, doi:10.1016/j.jmb.2018.02.007 (2018).

28. Ghosh, D. K. & Ranjan, A. An IRES-dependent translation of HYPK mRNA generates a truncated isoform of the protein that lacks the nuclear localization and functional ability. RNA Biol 16, 1604–1621, doi:10.1080/15476286.2019.1650612 (2019).

29. Ghosh, D. K., Roy, A. & Ranjan, A. Disordered Nanostructure in Huntingtin Interacting Protein K Acts as a Stabilizing Switch To Prevent Protein Aggregation. Biochemistry 57, 2009–2023, doi:10.1021/acs.biochem.7b00776 (2018).

30. Ghosh, D. K., Kumar, A. & Ranjan, A. Metastable states of HYPK-UBA domain’s seeds drive the dynamics of its own aggregation. Biochim Biophys Acta Gen Subj 1862, 2846–2861, doi:10.1016/j.bbagen.2018.09.003 (2018).

31. Arnesen, T. et al. The chaperone-like protein HYPK acts together with NatA in cotranslational N-terminal acetylation and prevention of Huntingtin aggregation. Mol Cell Biol 30, 1898–1909, doi:10.1128/MCB.01199-09 (2010).

32. Ayyadevara, S. et al. Proteins in aggregates functionally impact multiple neurodegenerative disease models by forming proteasome-blocking complexes. Aging Cell 14, 35–48, doi:10.1111/acel.12296 (2015).

33. Bell, R. et al. A human protein interaction network shows conservation of aging processes between human and invertebrate species. PLoS Genet 5, e1000414, doi:10.1371/journal.pgen.1000414 (2009).

34. Xu, G. et al. Vulnerability of newly synthesized proteins to proteostasis stress. J Cell Sci 129, 1892–1901, doi:10.1242/jcs.176479 (2016).

35. Azzam, M. E. & Algranati, I. D. Mechanism of puromycin action: fate of ribosomes after release of nascent protein chains from polysomes. Proc Natl Acad Sci U S A 70, 3866–3869, doi:10.1073/pnas.70.12.3866 (1973).

36. Yewdell, J. W., Anton, L. C. & Bennink, J. R. Defective ribosomal products (DRiPs): a major source of antigenic peptides for MHC class I molecules? J Immunol 157, 1823–1826 (1996).

37. Wenger, T. et al. Autophagy inhibition promotes defective neosynthesized proteins storage in ALIS, and induces redirection toward proteasome processing and MHCI-restricted presentation. Autophagy 8, 350–363, doi:10.4161/auto.18806 (2012).

38. Ciuffa, R. et al. The selective autophagy receptor p62 forms a flexible filamentous helical scaffold. Cell Rep 11, 748–758, doi:10.1016/j.celrep.2015.03.062 (2015).

39. Lu, B. et al. Identification of NUB1 as a suppressor of mutant Huntington toxicity via enhanced protein clearance. Nat Neurosci 16, 562–570, doi:10.1038/nn.3367 (2013).

40. Schulman, B. A. & Harper, J. W. Ubiquitin-like protein activation by E1 enzymes: the apex for downstream signalling pathways. Nat Rev Mol Cell Biol 10, 319–331, doi:10.1038/nrm2673 (2009).

41. Leidecker, O., Matic, I., Mahata, B., Pion, E. & Xirodimas, D. P. The ubiquitin E1 enzyme Ube1 mediates NEDD8 activation under diverse stress conditions. Cell Cycle 11, 1142–1150, doi:10.4161/cc.11.6.19559 (2012).

42. Liu, S. et al. NEDD8 ultimate buster-1 long (NUB1L) protein promotes transfer of NEDD8 to proteasome for degradation through the P97UFD1/NPL4 complex. J Biol Chem 288, 31339–31349, doi:10.1074/jbc.M113.484816 (2013).

43. Nedelsky, N. B., Todd, P. K. & Taylor, J. P. Autophagy and the ubiquitin-proteasome system: collaborators in neuroprotection. Biochim Biophys Acta 1782, 691–699, doi:10.1016/j.bbadis.2008.10.002 (2008).

44. Su, V. & Lau, A. F. Ubiquitin-like and ubiquitin-associated domain proteins: significance in proteasomal degradation. Cell Mol Life Sci 66, 2819–2833, doi:10.1007/s00018-009-0048-9 (2009).

45. Tanaka, T., Kawashima, H., Yeh, E. T. & Kamitani, T. Regulation of the NEDD8 conjugation system by a splicing variant, NUB1L. J Biol Chem 278, 32905–32913, doi:10.1074/jbc.M212057200 (2003).

46. Gottlieb, L. & Marmorstein, R. Structure of Human NatA and Its Regulation by the Huntingtin Interacting Protein HYPK. Structure 26, 925–935 e928, doi:10.1016/j.str.2018.04.003 (2018).

47. Habisov, S. et al. Structural and Functional Analysis of a Novel Interaction Motif within UFM1-activating Enzyme 5 (UBA5) Required for Binding to Ubiquitin-like Proteins and Ufmylation. J Biol Chem 291, 9025–9041, doi:10.1074/jbc.M116.715474 (2016).

48. Macharia, M. W., Tan, W. Y. Z., Das, P. P., Naqvi, N. I. & Wong, S. M. Proximity-dependent biotinylation screening identifies NbHYPK as a novel interacting partner of ATG8 in plants. BMC Plant Biol 19, 326, doi:10.1186/s12870-019-1930-8 (2019).

49. Keuss, M. J. et al. Unanchored tri-NEDD8 inhibits PARP-1 to protect from oxidative stress-induced cell death. EMBO J 38, doi:10.15252/embj.2018100024 (2019).

50. Zhang, X. et al. Hepatic neddylation targets and stabilizes electron transfer flavoproteins to facilitate fatty acid beta-oxidation. Proc Natl Acad Sci U S A 117, 2473–2483, doi:10.1073/pnas.1910765117 (2020).

51. Xie, P. et al. The covalent modifier Nedd8 is critical for the activation of Smurf1 ubiquitin ligase in tumorigenesis. Nat Commun 5, 3733, doi:10.1038/ncomms4733 (2014).

52. Sundqvist, A., Liu, G., Mirsaliotis, A. & Xirodimas, D. P. Regulation of nucleolar signalling to p53 through NEDDylation of L11. EMBO Rep 10, 1132–1139, doi:10.1038/embor.2009.178 (2009).

53. Gong, L. & Yeh, E. T. Identification of the activating and conjugating enzymes of the NEDD8 conjugation pathway. J Biol Chem 274, 12036–12042, doi:10.1074/jbc.274.17.12036 (1999).

54. Lobato-Gil, S. et al. Proteome-wide identification of NEDD8 modification sites reveals distinct proteomes for canonical and atypical NEDDylation. Cell Rep 34, 108635, doi:10.1016/j.celrep.2020.108635 (2021).

55. Johansen, T. & Lamark, T. Selective Autophagy: ATG8 Family Proteins, LIR Motifs and Cargo Receptors. J Mol Biol 432, 80–103, doi:10.1016/j.jmb.2019.07.016 (2020).

56. Mandell, M. A., Kimura, T., Jain, A., Johansen, T. & Deretic, V. TRIM proteins regulate autophagy: TRIM5 is a selective autophagy receptor mediating HIV-1 restriction. Autophagy 10, 2387–2388, doi:10.4161/15548627.2014.984278 (2014).

57. Yamamoto, A. & Simonsen, A. The elimination of accumulated and aggregated proteins: a role for aggrephagy in neurodegeneration. Neurobiol Dis 43, 17–28, doi:10.1016/j.nbd.2010.08.015 (2011).

58. Richet, E. et al. NUB1 modulation of GSK3beta reduces tau aggregation. Hum Mol Genet 21, 5254–5267, doi:10.1093/hmg/dds376 (2012).

59. Tanji, K. et al. Immunohistochemical localization of NUB1, a synphilin-1-binding protein, in neurodegenerative disorders. Acta Neuropathol 114, 365–371, doi:10.1007/s00401-007-0238-1 (2007).

60. Hodges, A. et al. Brain gene expression correlates with changes in behavior in the R6/1 mouse model of Huntington’s disease. Genes Brain Behav 7, 288–299, doi:10.1111/j.1601-183X.2007.00350.x (2008).

61. Raychaudhuri, S., Sinha, M., Mukhopadhyay, D. & Bhattacharyya, N. P. HYPK, a Huntingtin interacting protein, reduces aggregates and apoptosis induced by N-terminal Huntingtin with 40 glutamines in Neuro2a cells and exhibits chaperone-like activity. Hum Mol Genet 17, 240–255, doi:10.1093/hmg/ddm301 (2008).

62. Choudhury, K. R. & Bhattacharyya, N. P. Chaperone protein HYPK interacts with the first 17 amino acid region of Huntingtin and modulates mutant HTT-mediated aggregation and cytotoxicity. Biochem Biophys Res Commun 456, 66–73, doi:10.1016/j.bbrc.2014.11.035 (2015).

63. Sakurai, H., Sawai, M., Ishikawa, Y., Ota, A. & Kawahara, E. Heat shock transcription factor HSF1 regulates the expression of the Huntingtin-interacting protein HYPK. Biochim Biophys Acta 1840, 1181–1187, doi:10.1016/j.bbagen.2013.12.006 (2014).

64. Weyer, F. A. et al. Structural basis of HypK regulating N-terminal acetylation by the NatA complex. Nat Commun 8, 15726, doi:10.1038/ncomms15726 (2017).

65. Scott, D. C. et al. Blocking an N-terminal acetylation-dependent protein interaction inhibits an E3 ligase. Nat Chem Biol 13, 850–857, doi:10.1038/nchembio.2386 (2017)

66. Ghosh, D. K., Roy, A. & Ranjan, A. The ATPase VCP/p97 functions as a disaggregase against toxic Huntingtin-exon1 aggregates. FEBS Lett 592, 2680–2692, doi:10.1002/1873-3468.13213 (2018).

67. Kumar, A., Ghosh, D. K., Ali, J. & Ranjan, A. Characterization of Lipid Binding Properties of Plasmodium falciparum Acyl-Coenzyme A Binding Proteins and Their Competitive Inhibition by Mefloquine. ACS Chem Biol 14, 901–915, doi:10.1021/acschembio.9b00003 (2019).

68. Rueden, C. T. et al. ImageJ2: ImageJ for the next generation of scientific image data. BMC Bioinformatics 18, 529, doi:10.1186/s12859-017-1934-z (2017).

69. Loos, B., du Toit, A. & Hofmeyr, J. H. Defining and measuring autophagosome flux-concept and reality. Autophagy 10, 2087–2096, doi:10.4161/15548627.2014.973338 (2014).

70. Berman, H. M. et al. The Protein Data Bank. Nucleic Acids Res 28, 235–242, doi:10.1093/nar/28.1.235 (2000).

71. Ghosh, D.K., Shrikondawar, A.N. & Ranjan A. Local structural unfolding at the edge-strands of beta sheets is the molecular basis for instability and aggregation of G85R and G93A mutants of Superoxide dismutase 1. J Biomol Struct Dyn 38(3), 647–659, doi:10.1080/07391102.2019.1584125 (2019).

72. Ghosh, D.K., Kumar, A. & Ranjan A. T54R mutation destabilizes the dimer of Superoxide dismutase T54R by inducing steric clashes at the dimer interface. RSC Advances 10, 10776–10788, DOI: 10.1039/C9RA09870D (2020).

73. Hinsen, K. The Molecular Modeling Toolkit: A New Approach to Molecular Simulations. J. Comput. Chem., 21(2), 79–85, doi:10.1002/(SICI)1096-987X(20000130) (2000).

74. Case, D. A. et al. The Amber biomolecular simulation programs. J Comput Chem 26, 1668–1688, doi:10.1002/jcc.20290 (2005).

75. Kozakov, D., Brenke, R., Comeau, S. R. & Vajda, S. PIPER: an FFT-based protein docking program with pairwise potentials. Proteins 65, 392–406, doi:10.1002/prot.21117 (2006).

76. Krissinel, E. & Henrick, K. Inference of macromolecular assemblies from crystalline state. J Mol Biol 372, 774–797, doi:10.1016/j.jmb.2007.05.022 (2007).

77. Kozlov, G. et al. Structural basis for UBA-mediated dimerization of c-Cbl ubiquitin ligase. J Biol Chem 282, 27547–27555, doi:10.1074/jbc.M703333200 (2007).

78. Zhou, Z. R. et al. Differential ubiquitin binding of the UBA domains from human c-Cbl and Cbl-b: NMR structural and biochemical insights. Protein Sci 17, 1805–1814, doi:10.1110/ps.036384.108 (2008).

79. Sander, B., Xu, W., Eilers, M., Popov, N. & Lorenz, S. A conformational switch regulates the ubiquitin ligase HUWE1. Elife 6, doi:10.7554/eLife.21036 (2017).

80. Marx, A. et al. Structural variations in the catalytic and ubiquitin-associated domains of microtubule-associated protein/microtubule affinity regulating kinase (MARK) 1 and MARK2. J Biol Chem 281, 27586–27599, doi:10.1074/jbc.M604865200 (2006).

81. Panneerselvam, S., Marx, A., Mandelkow, E. M. & Mandelkow, E. Structure of the catalytic and ubiquitin-associated domains of the protein kinase MARK/Par-1. Structure 14, 173–183, doi:10.1016/j.str.2005.09.022 (2006).

82. Murphy, J. M. et al. Conformational instability of the MARK3 UBA domain compromises ubiquitin recognition and promotes interaction with the adjacent kinase domain. Proc Natl Acad Sci U S A 104, 14336–14341, doi:10.1073/pnas.0703012104 (2007).

83. Walinda, E. et al. Solution structure of the ubiquitin-associated (UBA) domain of human autophagy receptor NBR1 and its interaction with ubiquitin and polyubiquitin. J Biol Chem 289, 13890–13902, doi:10.1074/jbc.M114.555441 (2014).

84. Whitby, F. G., Xia, G., Pickart, C. M. & Hill, C. P. Crystal structure of the human ubiquitin-like protein NEDD8 and interactions with ubiquitin pathway enzymes. J Biol Chem 273, 34983–34991, doi:10.1074/jbc.273.52.34983 (1998).

85. Withers-Ward, E. S., Mueller, T. D., Chen, I. S. & Feigon, J. Biochemical and structural analysis of the interaction between the UBA(2) domain of the DNA repair protein HHR23A and HIV-1 Vpr. Biochemistry 39, 14103–14112, doi:10.1021/bi0017071 (2000).

86. Ryu, K. S. et al. Binding surface mapping of intra- and interdomain interactions among hHR23B, ubiquitin, and polyubiquitin binding site 2 of S5a. J Biol Chem 278, 36621–36627, doi:10.1074/jbc.M304628200 (2003).

87. Evans, C. L., Long, J. E., Gallagher, T. R., Hirst, J. D. & Searle, M. S. Conformation and dynamics of the three-helix bundle UBA domain of p62 from experiment and simulation. Proteins 71, 227–240, doi:10.1002/prot.21692 (2008).

88. Middleton, A. J. & Day, C. L. The molecular basis of lysine 48 ubiquitin chain synthesis by Ube2K. Sci Rep 5, 16793, doi:10.1038/srep16793 (2015).

89. Zhang, D., Raasi, S. & Fushman, D. Affinity makes the difference: nonselective interaction of the UBA domain of Ubiquilin-1 with monomeric ubiquitin and polyubiquitin chains. J Mol Biol 377, 162–180, doi:10.1016/j.jmb.2007.12.029 (2008).

90. Kieken, F., Spagnol, G., Su, V., Lau, A. F. & Sorgen, P. L. NMR structure note: UBA domain of CIP75. J Biomol NMR 46, 245–250, doi:10.1007/s10858-010-9397-9 (2010).

91. Chang, Y. G. et al. Solution structure of the ubiquitin-associated domain of human BMSC-UbP and its complex with ubiquitin. Protein Sci 15, 1248–1259, doi:10.1110/ps.051995006 (2006).

92. Ponting, C. P., Schultz, J., Milpetz, F. & Bork, P. SMART: identification and annotation of domains from signalling and extracellular protein sequences. Nucleic Acids Res 27, 229–232, doi:10.1093/nar/27.1.229 (1999).

93. Sievers, F. et al. Fast, scalable generation of high-quality protein multiple sequence alignments using Clustal Omega. Mol Syst Biol 7, 539, doi:10.1038/msb.2011.75 (2011).

94. Lesnick, T. G. et al. A genomic pathway approach to a complex disease: axon guidance and Parkinson disease. PLoS Genet 3, e98, doi:10.1371/journal.pgen.0030098 (2007).

95. Dunckley, T. et al. Gene expression correlates of neurofibrillary tangles in Alzheimer’s disease. Neurobiol Aging 27, 1359–1371, doi:10.1016/j.neurobiolaging.2005.08.013 (2006).

96. Lee, J. M. et al. A novel approach to investigate tissue-specific trinucleotide repeat instability. BMC Syst Biol 4, 29, doi:10.1186/1752-0509-4-29 (2010).

97. Ferraiuolo, L. et al. Microarray analysis of the cellular pathways involved in the adaptation to and progression of motor neuron injury in the SOD1 G93A mouse model of familial ALS. J Neurosci 27, 9201–9219, doi:10.1523/JNEUROSCI.1470-07.2007 (2007).

98. Chen-Plotkin, A. S. et al. Variations in the progranulin gene affect global gene expression in frontotemporal lobar degeneration. Hum Mol Genet 17, 1349–1362, doi:10.1093/hmg/ddn023 (2008).

99. Aydin, D. et al. Comparative transcriptome profiling of amyloid precursor protein family members in the adult cortex. BMC Genomics 12, 160, doi:10.1186/1471-2164-12-160 (2011).

100. Gatchel, J. R. et al. The insulin-like growth factor pathway is altered in spinocerebellar ataxia type 1 and type 7. Proc Natl Acad Sci U S A 105, 1291–1296, doi:10.1073/pnas.0711257105 (2008).

101. Han, M. H. et al. Janus-like opposing roles of CD47 in autoimmune brain inflammation in humans and mice. J Exp Med 209, 1325–1334, doi:10.1084/jem.20101974 (2012).

102. Lockstone, H. E. et al. Gene expression profiling in the adult Down syndrome brain. Genomics 90, 647–660, doi:10.1016/j.ygeno.2007.08.005 (2007).

103. Kim, M. Y. et al. hsp70 and a novel axis of type I interferon-dependent antiviral immunity in the measles virus-infected brain. J Virol 87, 998–1009, doi:10.1128/JVI.02710-12 (2013).

104. Gersten, M. et al. An integrated systems analysis implicates EGR1 downregulation in simian immunodeficiency virus encephalitis-induced neural dysfunction. J Neurosci 29, 12467–12476, doi:10.1523/JNEUROSCI.3180-09.2009 (2009).

105. Hur, J. et al. The identification of gene expression profiles associated with progression of human diabetic neuropathy. Brain 134, 3222–3235, doi:10.1093/brain/awr228 (2011).

